# Plasma membrane associated Receptor Like Kinases relocalise to plasmodesmata in response to osmotic stress

**DOI:** 10.1101/610881

**Authors:** Magali S. Grison, Philip Kirk, Marie Brault, Xu Na Wu, Waltraud X Schulze, Yoselin Benitez-Alfonso, Françoise Immel, Emmanuelle M. Bayer

**Author notes:** These authors equally contributed to the work (Philip Kirk, Marie Brault).

## Abstract

Plasmodesmata act as key elements in intercellular communication, coordinating processes related to plant growth, development and responses to environmental stresses. While many of the developmental, biotic and abiotic signals are primarily perceived at the plasma membrane (PM) by receptor proteins, plasmodesmata also cluster receptor-like activities and whether or not these two pathways interact is currently unknown.

Here we show that specific PM-located Leucine-Rich-Repeat Receptor-Like-Kinases (LRR-RLKs), KIN7 and IMK2, which under optimal growth conditions are absented from plasmodesmata, rapidly relocate and cluster to the pores in response to osmotic stress. This process is remarkably fast, it is not a general feature of PM-associated proteins and is independent of sterol- and sphingolipid-membrane composition. Focusing on KIN7, previously reported to be involved in stress responses, we show that relocalisation upon mannitol depends on KIN7 phosphorylation. Loss-of-function mutation in KIN7 induces delay in lateral root (LR) development and the mutant is affected in the root response to mannitol stress. Callose-mediated plasmodesmata regulation is known to regulate LR development. We found that callose levels are reduced in *kin7* mutant background with a root phenotype resembling ectopic expression of PdBG1, an enzyme that degrades callose at the pores. Both the LR and callose phenotypes can be complemented by expression of KIN7 -wild-type and –phosphomimic variants but not by KIN7 phosphodead mutant which fails to relocalise at plasmodesmata. Together the data indicate that re-organisation of RLKs to plasmodesmata is important for the regulation of callose and LR development as part of the plant response to osmotic stress.

## Introduction

Plasmodesmata are nano-scaled membranous pores that span the plant cell wall creating both cytoplasmic and membrane continuums between cells (Tilsner et al., 2016, 2011). By interconnecting most cells throughout the whole plant body, plasmodesmata form a symplastic network which supports and controls the movement of molecules from cell-to-cell, within a given tissue or organ, and the long-distance transport when combined with the vasculature (Corbesier, 2009; Kragler et al., 1998; Liu et al., 2012; Reagan et al., 2018). Given their central function in intercellular communication, plasmodesmata orchestrate processes related to plant growth and development but also responses to pathogens and abiotic stresses (Benitez-Alfonso et al., 2013, 2010; Caillaud et al., 2014; Cui and Lee, 2016; Daum et al., 2014; Faulkner et al., 2013; Gallagher et al., 2014; Lee et al., 2011; Lexy et al., 2018; Lim et al., 2016; Liu et al., 2012; Miyashima et al., 2019; Tylewicz and Bhalerao, 2018; Vaten et al., 2011; Wu et al., 2016). Plasmodesmata also act as specialised signalling hubs, capable of generating and/or relaying signalling from cell-to-cell through plasmodesmata-associated receptor–activity (Faulkner, 2013; Stahl et al., 2013; Stahl and Faulkner, 2015; Vaddepalli et al., 2014)

Plasmodesmata specialised functions hinges on their molecular specialisation (Bayer et al., 2004; Nicolas et al., 2017). The pores are outlined by highly-specialised plasma membrane microdomains which cluster a specific set of both proteins and lipids, compared to the bulk PM (Benitez-Alfonso et al., 2013; Fernandez-Calvino et al., 2011; Grison et al., 2015; Levy et al., 2007; Salmon and Bayer, 2013; Simpson et al., 2009; Thomas et al., 2008; Vaten et al., 2011; Xu et al., 2017). Amongst the array of proteins that localise to plasmodesmata, receptor proteins and receptor protein kinases have recently emerged as critical players for modulating cell-to-cell signalling in response to both developmental and stress-related stimuli (Faulkner et al., 2013; Stahl and Faulkner, 2015; Stahl and Simon, 2013; Vaddepalli et al., 2014). For instance, Plasmodesmata Located Protein 5 (PDLP5), a receptor-like protein, is necessary for callose induced-plasmodesmata closure in response to salicylic acid, a pivotal hormone in innate immune responses (Lee et al., 2011; Wang et al., 2013). Similarly, up-regulation of PDLP1 during mildew infection promotes down-regulation of plasmodesmata permeability (Caillaud et al., 2014). Membrane associated Receptor Like Kinases (RLKs), such as STRUBBELIG localises at plasmodesmata where it interacts with QUIRKY to regulate organ formation and tissue morphogenesis (Vaddepalli et al., 2014). Similarly, the receptor kinase CRINKLY4 presents dual localisation at the PM and plasmodesmata and is involved in root apical meristem maintenance and columella cell identity specification (Stahl et al., 2013). CRINKLY4 forms homo- and hetero-meric complexes with CLAVATA1, depending on its subcellular localisation at the PM or at plasmodesmata (Stahl et al., 2013). Activation/inactivation of signalling cascades often correlates with receptor complex association/dissociation to PM microdomains (Hofman et al., 2008). There is a high diversity of microdomains that co-exist at the PM allowing the separation of different signalling pathways (Bücherl et al., 2017; Jarsch et al., 2014; Raffaele et al., 2007). For instance in plants, the localisation of FLAGELLIN SENSING 2 and BRASSINOSTEROID INSENSITIVE 1 in distinct microdomains enable cells to differentiate between fungus-induced immunity and steroid-mediated growth, and this is despite the fact that these two signalling cascades share common components (Bücherl et al., 2017). In mammals, the EPIDERMAL GROWTH FACTOR RECEPTOR reversibly associates and dissociates with PM microdomains, which in turn control the activation and inactivation of signalling events (Bocharov et al., 2016; Hofman et al., 2008). Spatio-temporality and dynamics of receptor-complexes appears critical for regulating signalling events. In plants, both the PM and plasmodesmata pores present receptor-like activities but at present it is not clear whether these interact.

Here, we present data revealing that the PM-located Leucine Rich Repeat Receptor Like Kinases (LRR-RLKs), KIN7 (Kinase7; AT3G02880) and IMK2 (Inflorescence Meristem Kinase2; AT3G51740) rapidly re-organise their subcellular localisation and relocate at plasmodesmata intercellular pores, upon mannitol and NaCl treatments. This process occurs within less than 2 min and it is not a general behaviour of PM or microdomain-associated proteins. Focusing on KIN7, which has been previously shown to be involved in sucrose- and ABA-related responses and associated with lipid nanodomains (Isner et al., 2018; Szymanski et al., 2015; Wu et al., 2013), we show that relocalisation does not depend on sterol or sphingolipid membrane composition. KIN7 is phosphorylated in response to various abiotic stresses such as salt and mannitol-treatments (Chang et al., 2012; Chen et al., 2010; Hem et al., 2007; Hsu et al., 2009; Kline et al., 2010; Niittylä et al., 2007; Xue et al., 2013) and our data evidence that KIN7 phosphorylation is important for plasmodesmata localisation in control and mannitol-stress conditions. KIN7 phosphodead but not phosphomimic mutant is impaired in plasmodesmata localisation upon stress. Loss-of-function in KIN7 in *Arabidopsi*s results in a reduction in lateral root (LR) numbers in control conditions and affects root response to mannitol treatment. These phenotypes can be complemented by KIN7 wild-type protein and KIN7 phosphomimic, but not KIN7 phosphodead protein mutant. Our data further indicate that callose deposition at plasmodesmata is modified upon mannitol stress and that phosphorylation of KIN7 is important to regulate LR response to mannitol most likely *via* a mechanism that modulates the levels of callose.

The work emphasizes the dynamic nature of plasmodesmata membrane domains, which can within few minutes of stimulation recruit PM located receptor-like proteins that presumably trigger local mechanisms that regulate plasmodesmata aperture and, thereby, the developmental response to environmental stresses.

## RESULTS

### The PM-associated LRR-RLKs KIN7 and IMK2 dynamically associate with plasmodesmata in response to mannitol and salt treatments

A survey of the recently published *Arabidopsis* plasmodesmata-proteome (Brault et al., 2018) identified several members of the RLKs family present in the plasmodesmata fraction with clade III members being predominant (Supplemental Table. S1). As plasmodesmata have been reported to be composed of sterol- and sphingolipid-enriched microdomains (Grison et al., 2015; Nicolas et al., 2017), we focused on RLKs which may preferentially associate with lipid microdomains by cross-referencing the accessions with seven published Detergent Resistant Membrane (DRM) proteome (Demir et al., 2013; Keinath et al., 2010; Kierszniowska S, Seiwert B, 2009; Minami et al., 2009; Shahollari et al., 2005, 2004; Srivastava et al., 2013; Szymanski et al., 2015). By doing so, we identified two Leucine Rich Repeat (LRR) RLKs, Kinase7 (KIN7, AT3G02880) and Inflorescence Meristem Kinase 2 (IMK2, AT3G51740), which were relatively abundant in the plasmodesmata proteome and consistently identified in DRM fractions (Supplemental Table S2).

We next investigated the subcellular localisation of the two LRR-RLKs, by transiently expressing the proteins as green (GFP) fluorescent protein fusions in *Nicotiana Benthamiana* leaves followed by confocal imaging. Under control conditions, both KIN7 and IMK2 were found exclusively located to the PM with no specific enrichment at plasmodesmata (Fig. 1A-D). However, when subjected to 0.4 M Mannitol or 100 mM NaCl both proteins re-organise at the cell periphery in a punctate pattern (Fig. 1A, C arrows). Co-localisation with the plasmodesmata marker, PDLP1-mRFP (Amari et al., 2010), revealed that the mannitol- and salt-induced peripheral dots co-localised with plasmodesmata (Fig. 1A, C). In order to quantify plasmodesmata depletion/enrichment under control and stress conditions, we measured the plasmodesmata index, called PD index, by calculating the fluorescence intensity ratio between plasmodesmata (green signal that co-localizes with PDLP1-mRFP) *versus* PM (see Methods and Supplemental Fig. S1). In control conditions both KIN7 and IMK2 displayed a PD Index below 1 (median value) indicating no specific enrichment at plasmodesmata compared to the PM. However, upon short-term (5-30 min) mannitol or NaCl treatment this value raised up to 1.5-2 (Fig. 1B, D), confirming plasmodesmata enrichment. In addition to clustering at plasmodesmata, we also observed a re-organisation of the LRR-RLK KIN7 within the PM plane into microdomains at the surface of epidermal cells (Fig. 1E), from which the proton pump ATPase PMA2 (Morsomme et al., 1998) was excluded.

**Figure 1.**
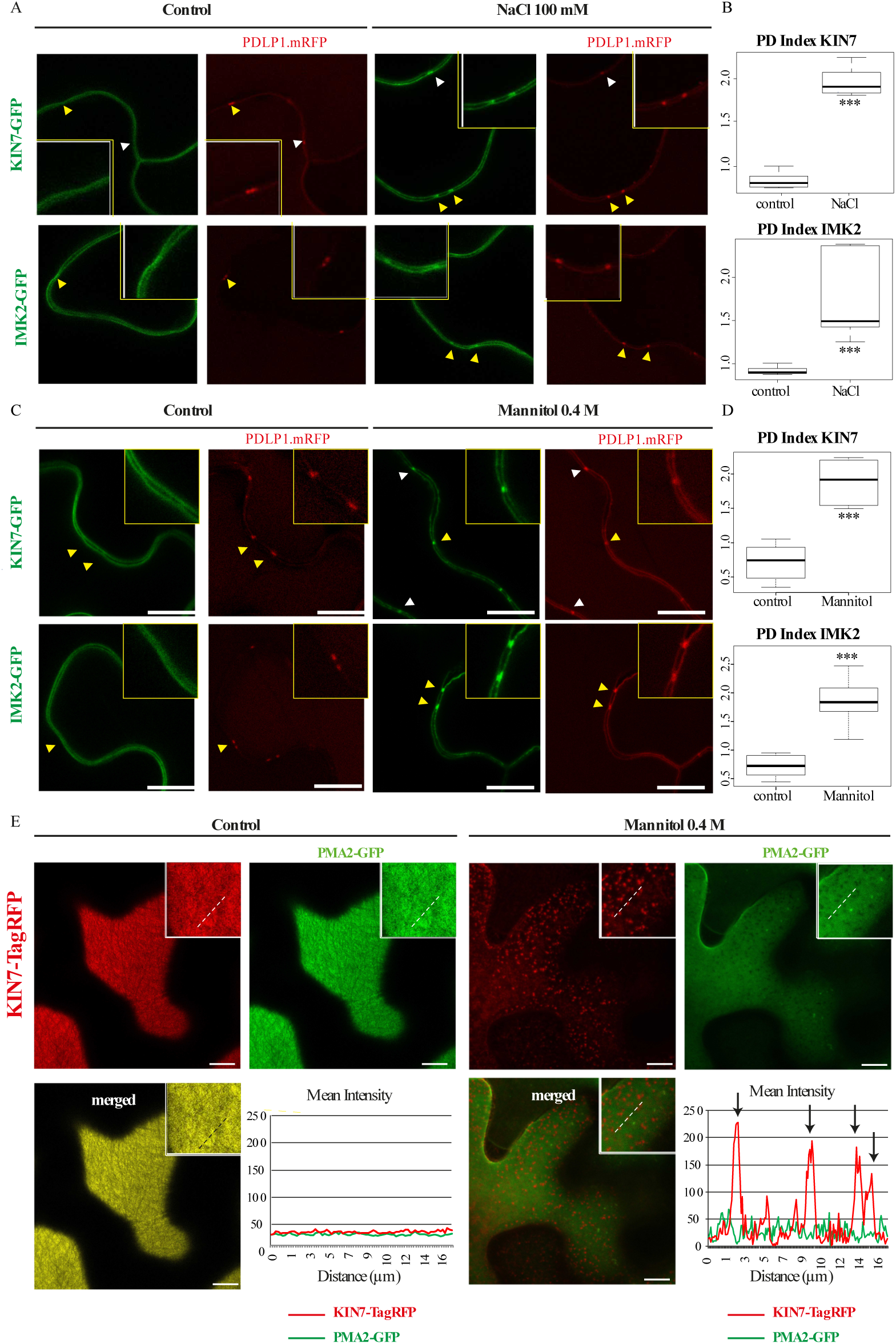
IMK2 and KIN7 are PM-associated LRR-RLKs that re-organise at plasmodesmata upon salt and mannitol treatments. A-D, Transient expression in *N. Benthamiana* epidermal cells of IMK2-GFP and KIN7-GFP LRR-RLKs expressed under 35S promoter and visualised by confocal microscopy. In control conditions, the two LRR-RLKs localise exclusively at the PM and present no enrichment at plasmodesmata, which are marked by PDLP1-mRFP. Upon NaCl 100 mM (A, B) or mannitol 0.4 M (C, D) treatment (5-30 min) the two LRR-RLKs relocalise to plasmodesmata (arrowheads). Yellow-boxed regions are magnification of areas indicated by yellow arrowheads. Enrichment at plasmodesmata versus the PM was quantified by the PD index, which correspond to the fluorescence intensity ratio of the LRR-RLKs at plasmodesmata versus the PM in control and abiotic stress conditions (see Methods for details and Supplemental Fig. S1). n=4 experiments, 3 plants/experiment, 10 measures/plant. Wilcoxon statistical analysis: * p-value <0.05; ** p-value<0.01; *** p-value <0.001 E, Transient expression in *N. Benthamiana* epidermal cells of KIN7-TagRFP and PMA2-GFP expressed under 35S promoter and visualised by confocal microscopy. Top surface view of a leaf epidermal cell showing the uniform and smooth distribution pattern of KIN7-TagRFP and PMA2-GFP at the PM under control conditions. Mannitol treatment causes a relocalisation of KIN7-TagRFP, but not of PMA2-GFP, into microdomain-like structures at the PM on the upper epidermal cell surface. Intensity plot along the white dashed line visible on the confocal images. n=2 experiments, 3 plants/experiment. Scale bars= 10µm.

To confirm these results, we generated *A. thaliana* transgenic lines expressing KIN7 tagged with GFP (Fig. 2). In control condition KIN7 was located to the PM in both cotyledons and root tissues of one week-old seedlings, but re-organised at the PM and relocated to plasmodesmata upon mannitol treatment (Fig. 2A-D). Re-organisation at plasmodesmata was remarkably fast and happened within 1 to 4 min post-treatment in the cotyledons (Fig. 2E; Supplemental Movie1). A similarly rapid change of localisation was also observed upon NaCl (100 mM) treatment (Supplemental Fig. S2).

**Figure 2.**
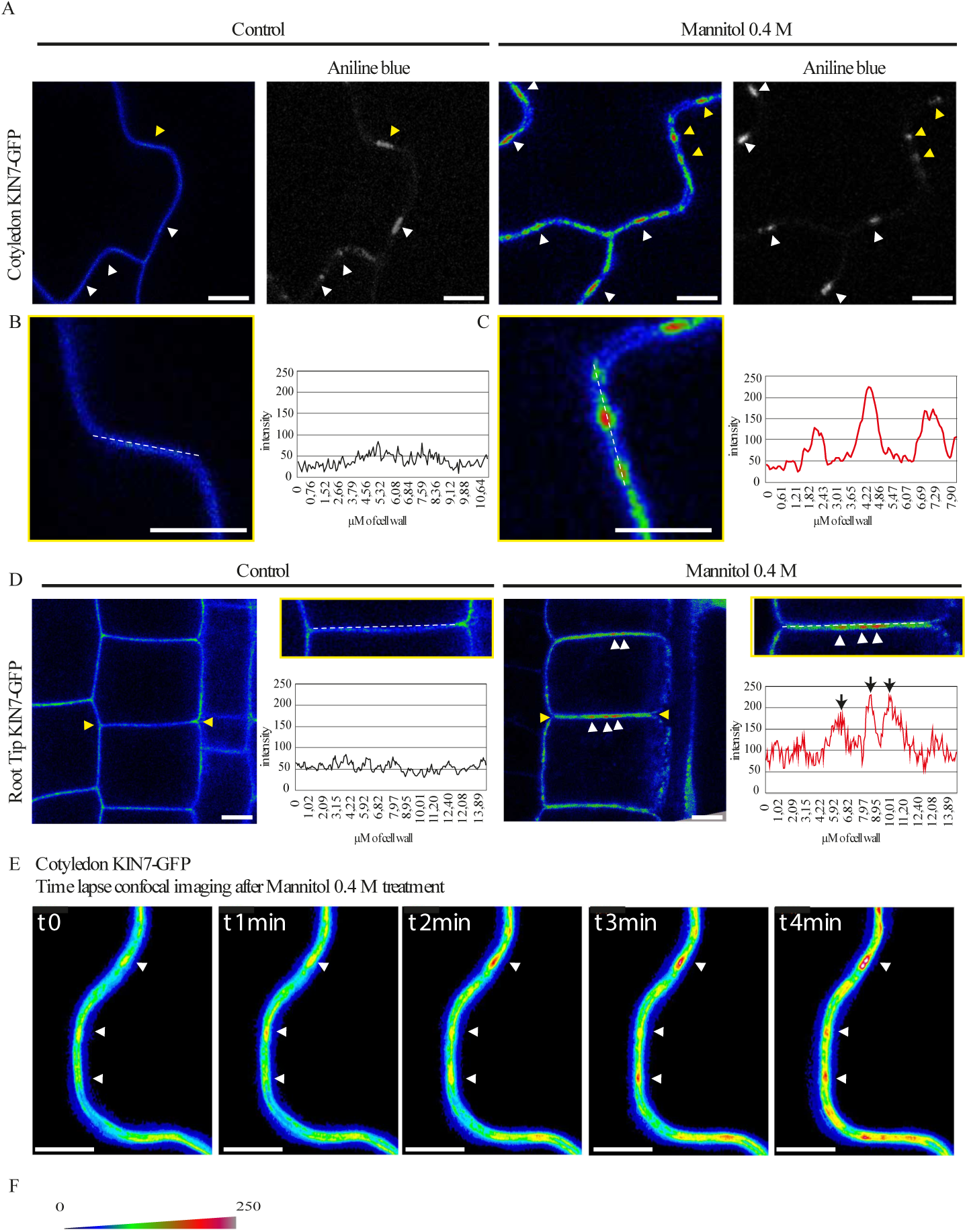
Re-organisation of KIN7 at plasmodesmata upon abiotic stress occurs remarquably fast. Stable *Arabidopsis* line expressing KIN7-GFP, under 35S promoter and visualised by confocal microscopy. All images have been color-coded through a heat-map filter to highlight clustering at plasmodesmata. A-D, In control conditions, KIN7-GFP localises exclusively at the PM in cotyledons (A-C) or root epidermis (D) and is not enriched at plasmodesmata (marked by aniline blue staining, arrowheads). B are magnified regions indicated by yellow arrowheads in A. Upon mannitol 0.4 M treatment, KIN7 relocalises to plasmodesmata where it becomes enriched (A and D, white arrowheads). Intensity plots along the white dashed lines are shown for KIN7-GFP localisation pattern in control and mannitol conditions. E, Time-lapse imaging of KIN7-GFP relocalisation upon mannitol exposure. Within less than two minutes plasmodesmata localisation already visible (white arrowhead). Please note re-organisation is faster when KIN7 is stably expressed (less than 5 min when stably expressed, 5-30 min when transiently expressed) F, Shows a color-coding bar for heat-map images. Scale bars= 10 μm

From our data we concluded that both KIN7 and IMK2 LRR-RLKs can rapidly modulate their subcellular localisation and associate with plasmodesmata in response to osmotic stress.

### Relocalisation at plasmodesmata is not a general feature of PM or nanodomain-associated proteins

To test whether plasmodesmata association in response to osmotic stress is a common feature of PM proteins, we investigated the behaviour of unrelated PM-associated proteins. We selected proteins that associate with the PM either through transmembrane domains, such as the Low Temperature Induced Protein 6B (Lti6b), the Plasma Membrane Intrinsic Protein 2;1 (PIP2;1) and PMA2 (Cutler et al., 2000; Prak et al., 2008), or through surface interaction with inner leaflet lipids such as Remorin 1.2 and 1.3, which are also well-established lipid nano-domain markers (Gronnier et al., 2017; Jarsch et al., 2014; Konrad et al., 2014). While KIN7 became significantly enriched at plasmodesmata, none of the tested PM-associated proteins displayed plasmodesmata association upon short (1-5 min) 0.4 M mannitol treatment as indicated by their PD index, which remained below 1 (Fig. 3A-B).

**Figure 3.**
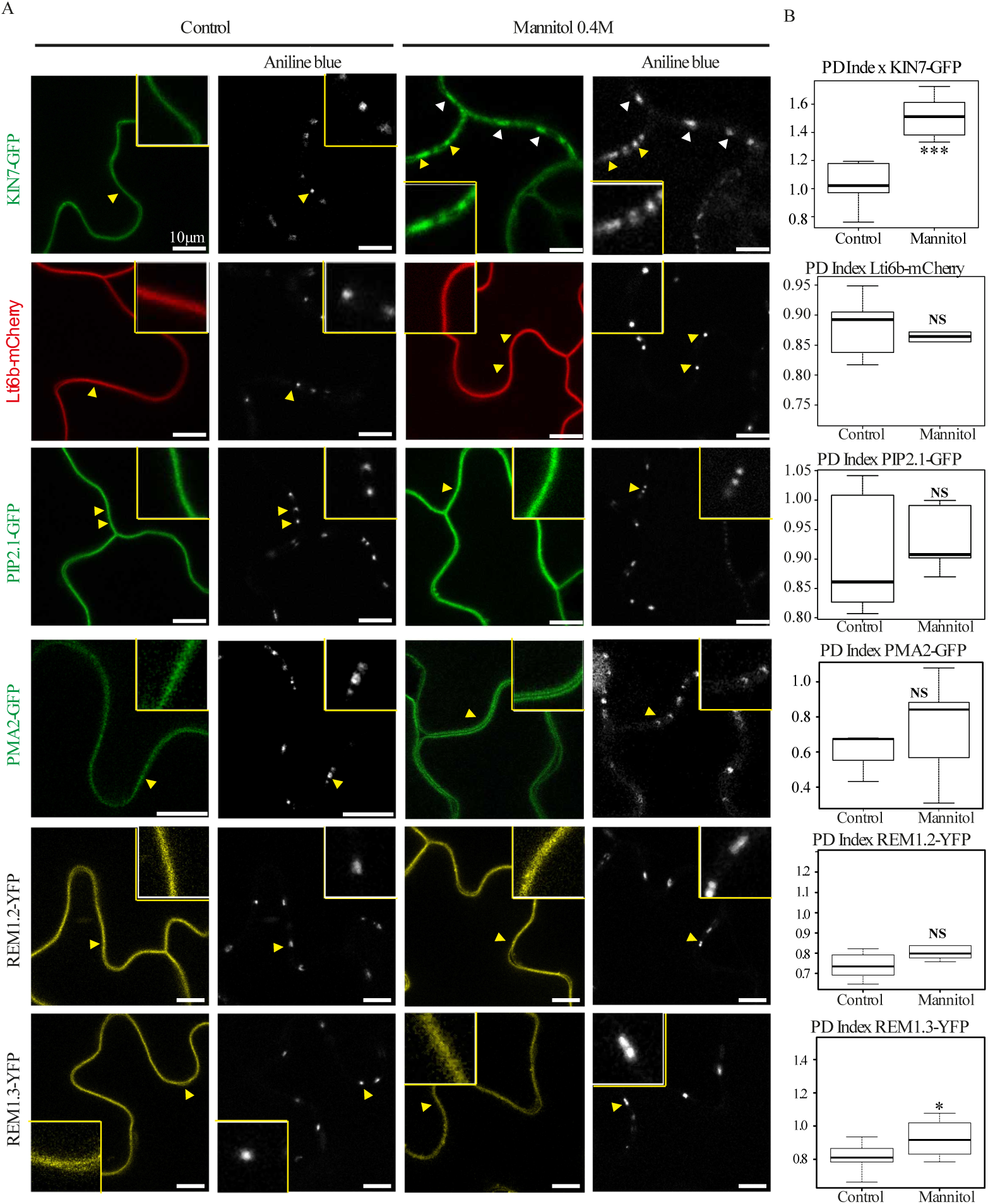
Conditional plasmodesmal association is not a general feature of PM-associated proteins. A, In control conditions, KIN7-GFP, the PM-associated protiens Lti6b-mCherry, PIP2;1-GFP, PMA2-GFP, REM1.2-YFP and REM1.3-YFP show localisation to the PM and are not enriched at plasmodesmata (stained with aniline blue, arrowheads). Mannitol 0.4 M treatment (1-5 min) induces the re-organisation of KIN7 at plasmodesmata, while other PM-associated proteins stay excluded from plasmodesmata. Single confocal scan images of *Arabidopsis* transgenic seedlings (KIN7-GFP, Lti6b-mCherry, PIP2;1-GFP, REM1.2-YFP and REM1.3-YFP) or *N. benthamiana* leaves transiently expressing PMA2-GFP. Yellow boxed regions are magnifications of areas indicated by yellow arrowheads. B, PD index for each PM-associated protein tested in A in control and mannitol conditions. n=3, 3 plants/line/experiment, 3 to 6 cells/plant, 5-10 ROI for PM and plasmodesmata per cell. Wilcoxon statistical analysis: * p-value <0.05; ** p-value<0.01; *** p-value <0.001. Scale bar=10µm

Altogether our results indicate that the capacity of KIN7 and IMK2 to relocalise at plasmodesmata upon stress is not a general feature of all PM proteins.

### Changes in sterols and sphingolipids composition do not affect KIN7 conditional association with plasmodesmata

We next decided to investigate the mechanisms underlying plasmodesmata localisation of LRR-RLKs by focusing on KIN7. KIN7 has been proposed to associate with sterol- and sphingolipid-enriched PM nano-domains in plants (Demir et al., 2013; Keinath et al., 2010; Kierszniowska S, Seiwert B, 2009; Minami et al., 2009; Shahollari et al., 2005, 2004; Srivastava et al., 2013; Szymanski et al., 2015) (Supplemental Table. S2), and in animal cells lipid-nano-domains have been reported to coalesce and form signalling platforms in a sterol-dependant manner (Gaus, 2014).

To test the importance of lipids, for plasmodesmal conditional association, we used pharmacological approaches and specifically inhibited sterols and sphingolipids biosynthesis (Grison et al., 2015; He et al., 2003; Wattelet-Boyer et al., 2016). For sterols, we used fenpropimorph (FEN100; 100 μg/mL, 48 h) which acts directly in the sterol biosynthetic pathway by inhibiting the cyclopropyl-sterol isomerase, and which effects are well characterized in *Arabidopsis* seedlings (Hartmann et al., 2002; He et al., 2003). For sphingolipids, we focused on Glycosyl-Inositol-Phospho-Ceramides (GIPCs) which are the main sphingolipids associated with both plasmodesmata and lipid nano-domains (Cacas et al., 2016; Grison et al., 2015). We modulated GIPCs content, using metazachlor (MZ100; 100 nM/mL, 48 h) which reduces the very long chain fatty acid and hydroxylated very long chain fatty acid (VLCFA>24C and hVLCFA>24C) of GIPCs (Wattelet-Boyer et al., 2016). Alteration of the cellular pool of sterols and VLCFA-derived GIPCs was confirmed by gas chromatography coupled to mass spectrometry (Fig. 4E,F). We observed a depletion of 22.6 % of sterols and 30 % of hVLCFA and VLCFA consistent with previous studies (Grison et al., 2015; Wattelet-Boyer et al., 2016). Effectiveness of lipid inhibitor treatments on the PM lipid pool was also confirmed by the change of Remorin 1.2 organisation at the PM surface from nano-domains to a smooth pattern (Fig. 4D).

**Figure 4.**
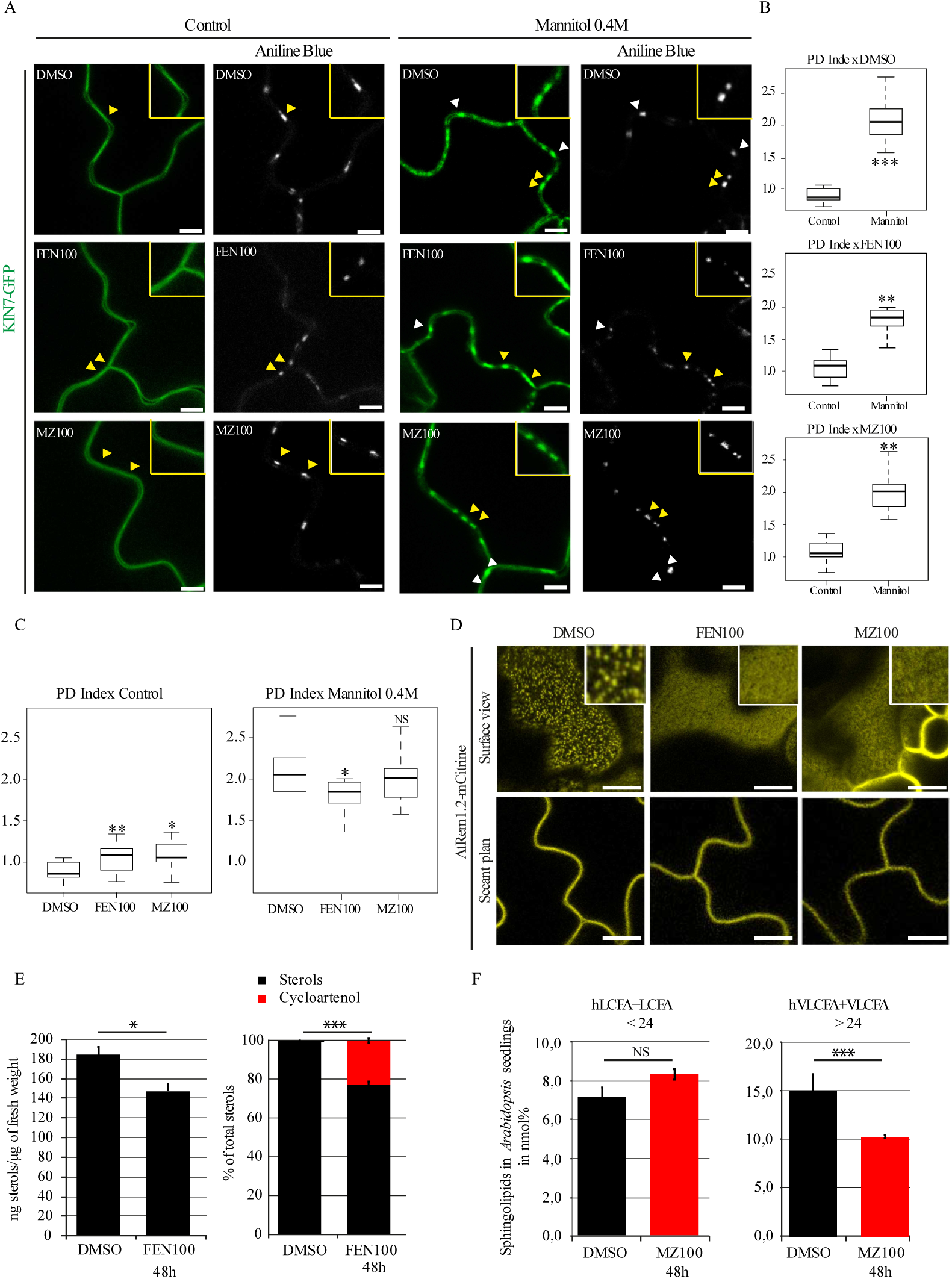
Mannitol-induced relocalisation of KIN7 is independent of sterols and sphingolipids. A-C, Stable *Arabidopsis* line expressing KIN7-GFP, under 35S promoter and visualised by confocal microscopy after sterol- or very long chain GIPC-biosynthesis inhibitor treatments and mannitol 0.4 M exposure (1-5min). Arabidopsis seedlings were grown on normal agar plates for 5 days and then transferred to 100 µg/mL Fenpropimorph (FEN100), 100 nM Metazachlor (MZ100) or 3% DMSO agar plates for 48h. Compared to control (DMSO) conditions, FEN100 and MZ100 induce a slight increase in plasmodesmata localisation as indicated by the PD index (B, C) but KIN7-GFP was still preferentially located at the PM. Despite the lipid inhibitor treatments KIN7-GFP was nevertheless capable of re-organising at plasmodesmata after mannitol treatment. A, Confocal single scan images. Yellow-boxed regions are magnification of areas indicated by yellow arrowheads. B, C, PD indexes corresponding to panel A. n=3 experiments, 3 plants/line/experiment, 3 to 6 cells/plant, 5-10 ROI for PM and plasmodesmata per cell. D, Localisation pattern of AtREM1.2-mCitrine in *Arabidopsis* cotyledons after 48h FEN100 and MZ100 treatments showing reduced lateral organisation into microdomains at the epidermal cell surface upon lipid inhibitors. E, Sterol quantification after FEN100 treatment by gaz chromatography coupled to mass spectrometry. Left, *Arabidopsis* seedlings treated with FEN100 presented a 20% decrease of the total amount of sterols after 48h. Right, relative proportion of sterol species in *Arabidopsis* seedling treated with FEN100 showing cycloartenol accumulation of 22,5%. Black: “normal” sterols; Red: cyloartenol. (n=3) Bars indicate SD. F, Total Fatty Acid Methyl Esthers (FAMES) quantification after MZ100 treatment by gaz chromatography coupled to mass spectrometry. VLCFA >24 (hydroxylated and non-hydroxylated) are reduced by 30% on metazaclhor. (n=3) Bars indicates SD. Wilcoxon statistical analysis: * p-value <0.05; ** p-value<0.01; *** p-value <0.001; **** p-value <0,0001. Scale bar= 10µm

Under conditions with no mannitol but sterol- and sphingolipid-inhibitors, we observed a minor but significant increase in the PD index of KIN7 under FEN100 and MZ100, which raised to 1.08 and 1.06, respectively, compared to DMSO control conditions with a PD index of 0.86 (Fig.4 C). The results indicate that modifying the cellular lipid pool can affect localisation to plasmodesmata. However, upon mannitol treatment (0.4 M, 1-5 min), effective KIN7 relocalisation to plasmodesmata was maintained in all conditions (Fig.4 A-C).

These results suggest that sterols and sphingolipids are not essential for plasmodesmata clustering of KIN7 under mannitol treatment.

### KIN7 association with plasmodesmata is regulated by phosphorylation

We next investigated whether KIN7 phosphorylation status could be involved in plasmodesmata targeting. Several phosphorylation sites have been experimentally reported for KIN7 (Supplemental Table. S3). KIN7 phospho-status varies upon various abiotic stresses such as salt and mannitol-treatments but also after exposure to sucrose and to hormones (Chang et al., 2012; Chen et al., 2010; Hem et al., 2007; Hsu et al., 2009; Kline et al., 2010; Niittylä et al., 2007; Xue et al., 2013). In the context of this study, we focused on two phosphorylation sites (S621 and S626), which were consistently and experimentally detected in several phosphoproteomic studies, including in response to salt and mannitol exposure (Supplemental Table. S3).

To test whether the phosphorylation of KIN7 could play a role in plasmodesmata association, we generated two KIN7 phosphomutants; the phosphomimic mutant (KIN7-S621D-S626D named hereafter KIN7-DD) and the phosphodead mutant (KIN7-S621A-S626A named hereafter KIN7-AA). Both were tagged with GFP, stably expressed under 35S in *Arabidopsis* and their localisation pattern analysed along with that of the wild type KIN7 protein (Fig. 5). Under control conditions, KIN7 and the phosphodead mutant KIN7-AA were localised at the PM (Fig.5A) and yielded PD indexes of 1.02 and 0.99 (median values; Fig. 5B-C), respectively indicating no specific plasmodesmata enrichment. By contrast KIN7-DD displayed a significantly higher PD index of 1.24 suggesting that, in control conditions, the phosphomimic mutant is already associated to plasmodesmata (Fig. 5A-C). Mannitol exposure (0.4 M Mannitol; 1-5 min treatment) triggered relocalisation of all proteins to a different extent. While KIN7 and KIN7-DD displayed a comparable PD index of 1.51 and 1.52 respectively, the phosphodead variant KIN7-AA, displayed a PD index barely reaching 1.20 (median values; Fig. 5B-C).

**Figure 5.**
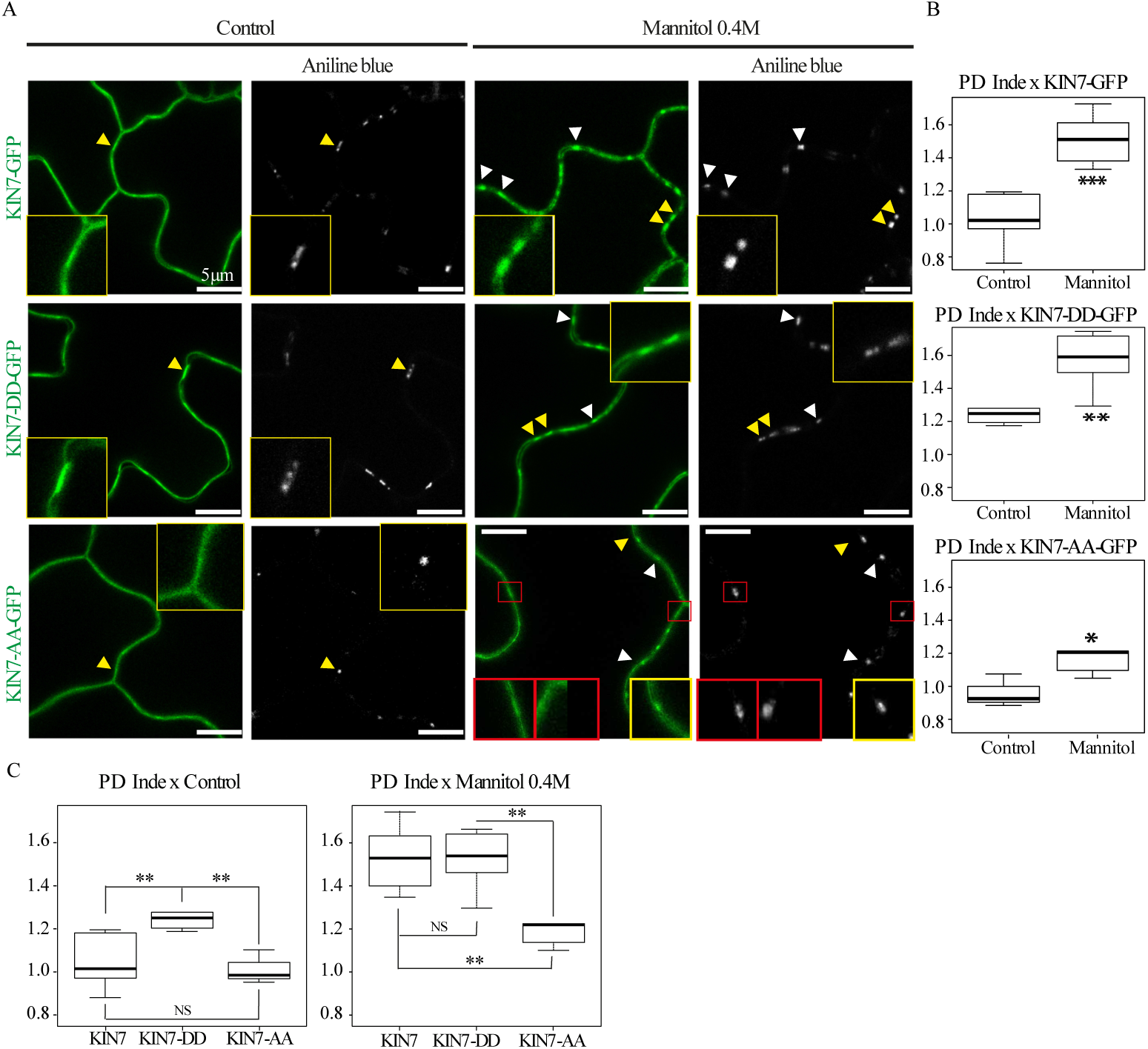
KIN7 phosphorylation regulates plasmodesmata association upon mannitol treatment. A-C, Stable *Arabidopsis* lines expressing KIN7-GFP, KIN7-DD-GFP (phosphomimic variant S621D-S626D) and KIN7-AA-GFP (phosphodead variant S621A-S626A) under 35S promoter and visualised by confocal microscopy. Plasmodesmata were labelled by aniline blue (arrowheads). In control condition KIN7 and the phosphodead mutant, KIN7-AA showed a “smooth” localisation pattern at the PM (A) with no significant plasmodesmata association (B, C). The phosphomimic KIN7-DD however, displayed a weak but significant plasmodesmata localisation with a shift of its PD index from 0.99 to 1.20 (A-C). After mannitol (0.4 M) exposure (1-5 min), KIN7 and KIN7-DD similarly relocalise at plasmodesmata with a PD index of 1.52 and 1.53, respectively. Re-organisation to plasmodesmata was significantly less effective for KIN7-AA (A-C), with a PD index barely reaching 1.20 upon mannitol. For the phosphodead KIN7-AA mutant, plasmodesmata-association was not systematic as shown in red boxes in A. A, Confocal single scan images. Yellow-boxed regions are magnification of areas indicated by yellow arrowheads. B, C PD indexes corresponding to panel A. n=3 experiments, 3 plants/line/experiments, 3 to 6 cells/plants, 5 to 10 ROI for PM and PD/cells. Wilcoxon statistical analysis: * p-value <0.05; ** p-value<0.01; *** p-value <0.001. Scale bars= 10µm.

From these data we concluded that KIN7 phosphorylation status influence plasmodesmata association and that mutations in the S621 and S626 phosphosites significantly alters KIN7 re-organisation at the pores.

### KIN7 function in modulating root development and response to mannitol

Osmotic stress and mannitol treatments are known to affect root system architecture (Deak et al., 2005; Kumar et al., 2019; MacGregor et al., 2008; Roycewicz and Malamy, 2012; Zhou et al., 2018). KIN7 localizes to plasmodesmata in response to mannitol and mutants in callose degradation and plasmodesmata transport are impaired in LR density and patterning (Benitez-Alfonso et al., 2013; Maule et al., 2013). We therefore tested KIN7 involvement in this pathway by determining its role in root development and in response to mannitol.

We first established the root phenotype of wild type Col-0 seedlings in mannitol (0.4M). After 3 days of exposure to mannitol, root length and LR number were reduced in comparison to seedlings in control media (Fig 6A-B). Mannitol treatment also modified callose, which appears reduced in internal root layers and increased in the epidermal cell layer (Fig. 7A-C) with a concomitant reduction of GFP symplastic movement into the epidermal cells when expressed under the SUC2 promoter (Fig. 7D-E).

**Figure 6.**
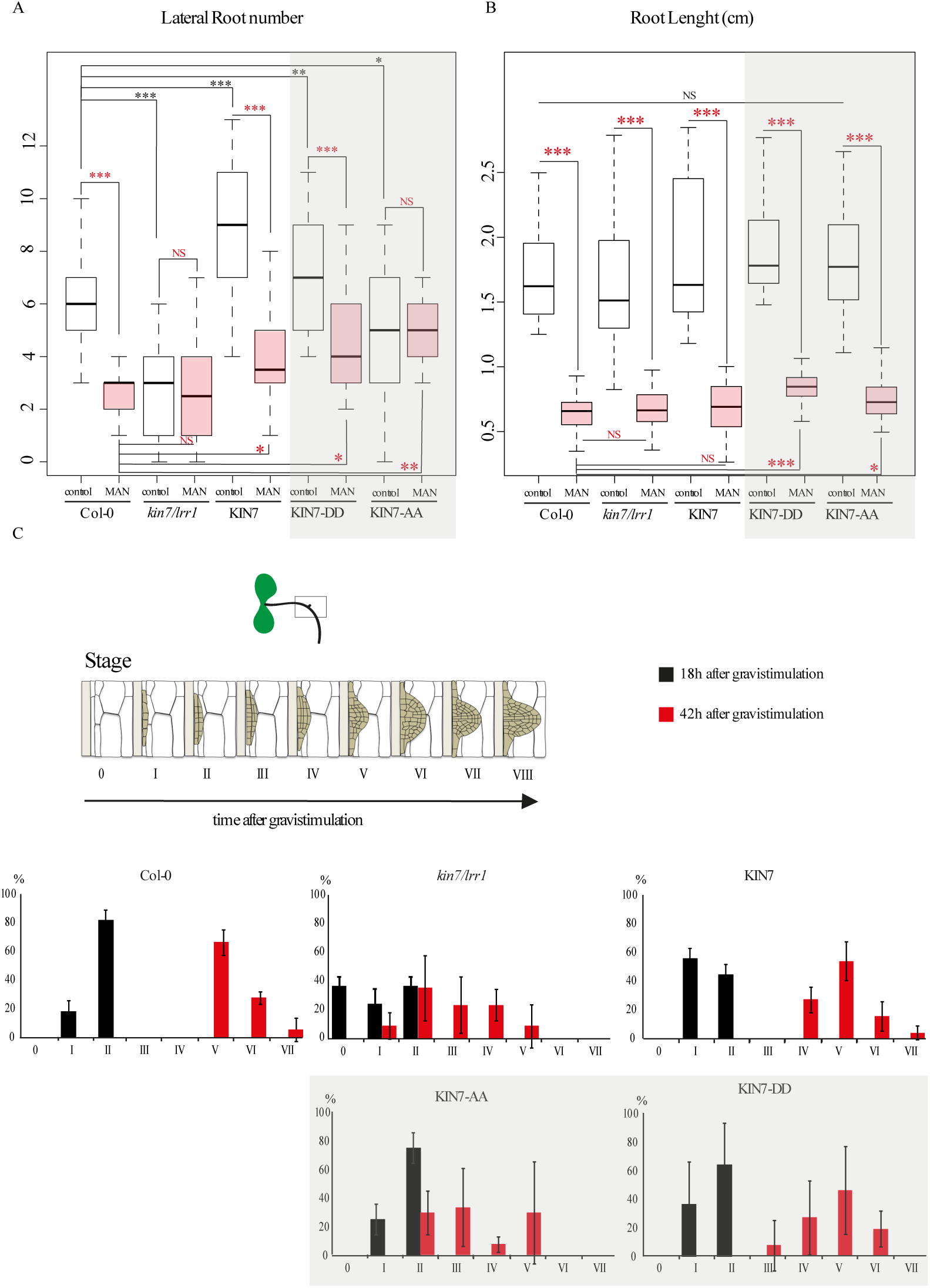
KIN7 is involved in root development and response to mannitol. A, LR number in wild type Col-0, kin7.*lrr1* mutant, kin7.*lrr1* expressing KIN7-GFP, KIN7-DD-GFP, KIN7-AA-GFP under 35S promoter. *Arabidopsis* lines were grown for 9 days on MS plates for control conditions, or 6 days then transferred to MS plate containing 0.4 M mannitol before root phenotyping. LR number is represented by white and red box plots for control and mannitol treatment, respectively. In control conditions, *kin7.lrr1* mutant displays a decrease of LR number compared to the wild type. Overexpression of KIN7 and the phosphomimic KIN7-DD reverse this phenotype with more LR. Overexpression of KIN7-AA phosphodead only partially rescues *kin7.lrr1* LR number phenotype. In response to mannitol treatment, Col-0 wild type and *Arabidopsis* seedlings overexpressing KIN7 and KIN7-DD in *kin7.lrr1* mutant background all showed a decrease in LR number, whereas *kin7.lrr1* and *kin7.lrr1* overexpressing KIN-AA display the same number of LR as in control conditions. B, The primary root length was measured in parallel to the LR (A) using FIJI software. None of the lines tested presented a significant root length difference compare to Col-0 in control conditions (white box plot). After mannitol treatment, all the lines were similarly affected with a reduction of the primary root length (red box plot), with the KIN7-DD and KIN7-AA showing a slight increase in their root length compared to Col-0. n=2 experiments, 10 plants/line/experiments. Wilcoxon statistical analysis: * p-value <0.05; ** p-value<0.01; *** p-value <0.001. Scale bars= 10µm. C, LR primordium stages, Top, Graphical summary of the gravistimulation and the development stages of the LR primordia adapted from Péret *et al.* 2012. Bottom, the LR primordium stages were determined 18h and 42h after gravistimulation, and are color-coded respectively in black and red. At 18h, the *kin7.lrr1* mutant display a delay in LR primordium initiation with the absence of LR primordium initiation (stage 0) in 35% of the plants observed. At 42h both the *kin7.lrr1* mutant and KIN7-AA-GFP expressing lines showed a delay in LR primordium compared to other lines, with no stage VI or VII LR primordium.

**Figure 7.**
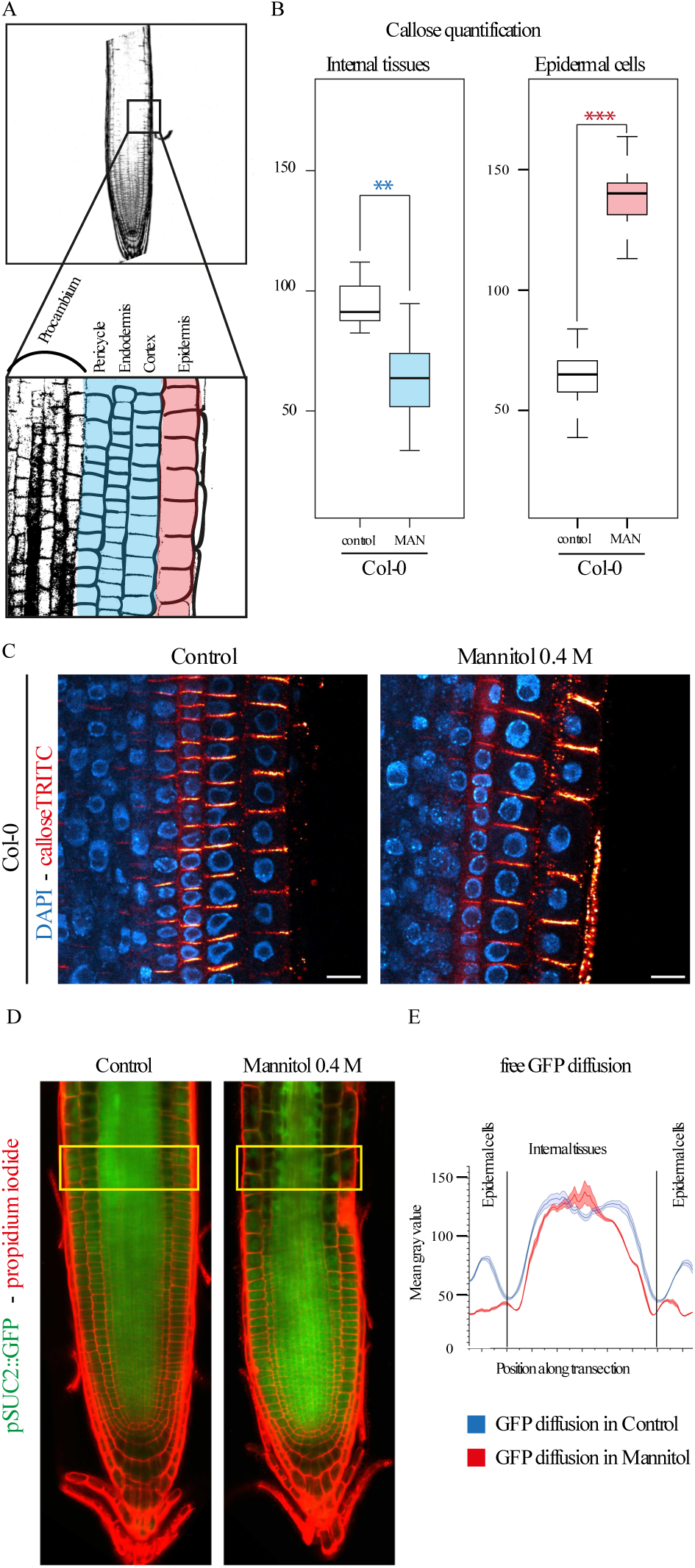
Callose and plasmodesmata trafficking is modulated upon mannitol treatment. A-C, A, representative scheme showing the root cell lineage with epidermal cells coloured in red and “internal layers” coloured in blue. The same colour code has been conserved in the box plot representation to facilitate the lecture of the figure. B, Callose level quantifications; upon mannitol treatment (3h, 0.4 M mannitol) callose levels are down regulated in internal layers (blue) of the root while being up regulated in the epidermis (red). C, Representative confocal images of callose immunofluorescence (red) in wild type Col-0 *Arabidopsis* roots in control and mannitol treatment. DAPI staining of DNA (blue) was performed to highlight the cellular organisation of root tissues. Scale bar 10 μm. D-E, *Arabidopsis* seedlings expressing pSUC2::GFP in under control and mannitol treatment (16h, 0.4 M mannitol). GFP symplastic unloading from the phloem to surrounding tissues is modified under mannitol treatment. We observed a reduction of GFP diffusion in epidermal cells, which showed increased callose levels at plasmodesmata (panels B-C). Scale bar 50 μm.

Next, we compared the root phenotype of the wild type Col-0 and loss-of-function KIN7 *Arabidopsis* mutant grown in parallel. Since KIN7 shares more than 90% similarity at the amino acid level to the LRR-RLK LRR1 (AT5G16590) and these proteins also display very similar expression profiles (Supplemental Fig. S3 and S4), we generated a double loss-of-function mutant named *kin7.lrr1.* The *kin7.lrr1* mutant and the overexpressor line 35S::KIN7-GFP in the mutant background (see Supplemental Fig. S5 for expression levels) were grown in MS control media and root phenotype was analysed 9 days after germination. We found that the primary root length was not significantly different between Col-0, *kin7.lrr1* and *link7.lrr1* overexpressing KIN7 (Fig.6B, white box plots). LR development, on the other hand, was significantly affected in the *kin7.lrr1* mutant and the KIN7 overexpressing line, with *kin7.lrr1* displaying a reduced number of LR and KIN7 over expressor showing the opposite phenotype with an increase in LR number in comparison to wild type (Fig.6 A, white box plots).

To further dissect this phenotype we examined the different stages of LR formation by subjecting the seedlings to a 90° gravitropic stimulus, which triggers LR initiation in a very synchronized manner at the outer edge of the bend root (Péret et al., 2012). LR initiation and outgrow was observed at 18h and 42h post-gravitropic stimuli (Fig. 6 C). LR initiation was impaired in the *kin7.lrr1 Arabidopsis* mutant as 35% of the bend roots did not display LR primordium 18h after gravistimulation and no stage VI and VII primordia were found after 42h. Over-expression of KIN7, on the other hand, resulted in only a slight delay in LR development. We also tested the response of the *kin7.lrr1* mutant and KIN7 overexpressing line to mannitol treatment. Mannitol caused a similar reduction in root length in all the lines tested, i.e. *kin7.lrr1*, KIN7 overexpressing seedlings and Col-0 wild type (Fig.6 A-B, compare white and red boxes). However, while Col-0 wild type showed reduced number of LR in mannitol compare to control growth conditions, *kin7.lrr1* was not significantly affected (Fig. 6A, compare white and red box plots). Hence, in *kin7.lrr1* mutant the number of LR was not reduced further by mannitol exposure in comparison to control growth conditions. Expression of KIN7 in *kin7.lrr1* background complemented the phenotype restoring LR response (reduced LR number) to mannitol (Fig. 6A). In summary, LR development and response to mannitol is significantly affected by mutation in KIN7.

Since mannitol induces changes in callose deposition (Fig.7), we used immunolocalization to detect callose levels in *kin7.lrr1* mutant and KIN7 overexpressor line (Fig. 8). The *kin7.lrr1* mutant showed reduced callose levels compared to wild type seedlings, while the over-expressing KIN7 lines appear to accumulate more callose (Fig.8A-B). These results suggest that callose down regulation may be accountable for *kin7.lrr1* LR phenotype. To test this hypothesis, we studied the root phenotype in a line ectopically expressing PdBG1, a plasmodesmata associated β1-3 glucanase (AT3G13560) which degrades callose (Benitez-Alfonso et al., 2013; Maule et al., 2013). Similarly to *kin7.lrr1* mutant, over-expression of PdBG1 did not affect primary root length PdBG1 but LR number was reduced compared to Col-0 in control conditions (Fig.8 C). After mannitol treatment changes in LR number were reduced in the PdBG1 overexpressor to a lesser extent than wild type, partially resembling *kin7.lrr1* response. This suggests that ectopic callose degradation is, at least partly, related to the LR response in control and mannitol growth conditions.

**Figure 8.**
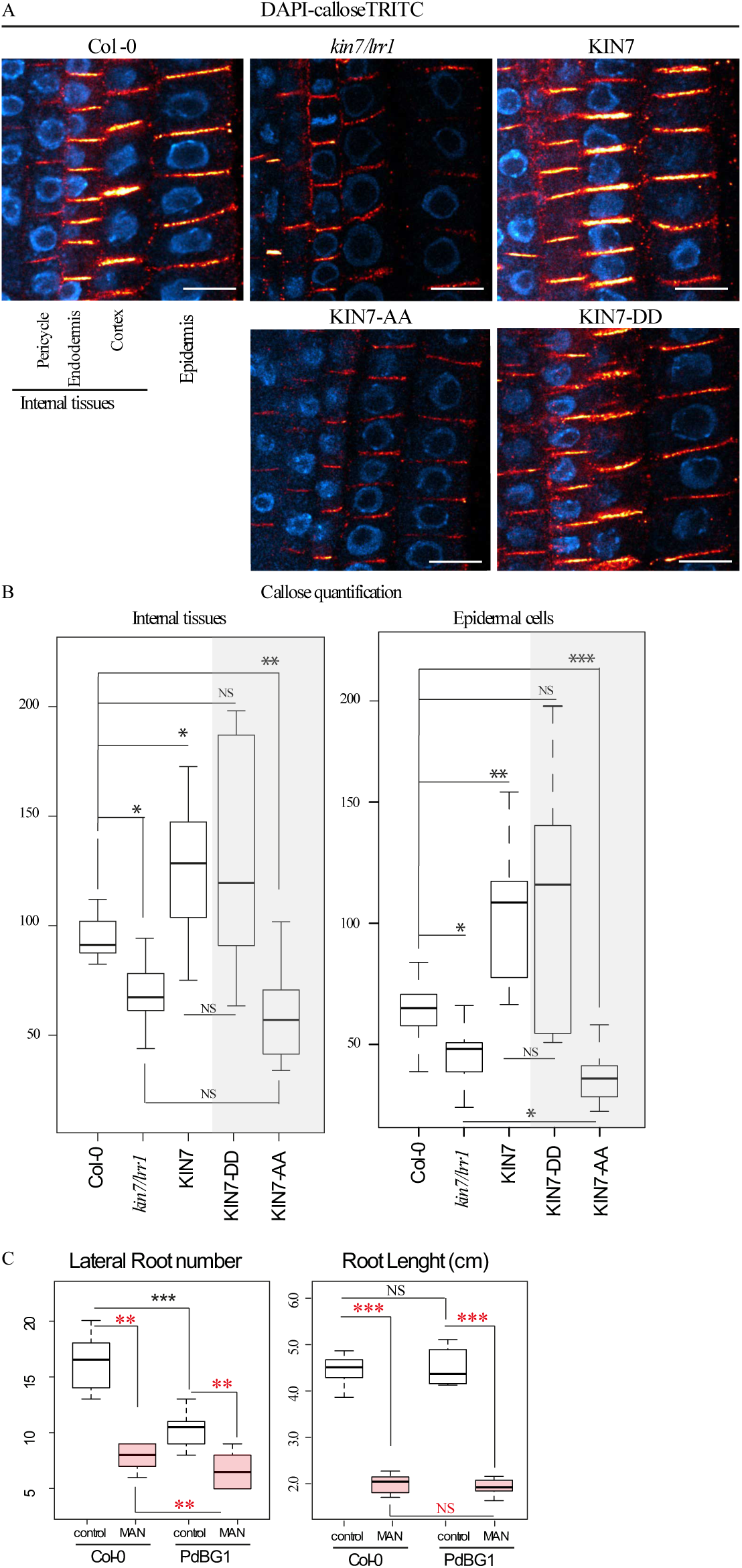
KIN7 is involved in callose regulation at plasmodesmata, which depends on KIN7 phosphorylation status. A-B, Quantification of callose levels in Col-0, *kin7.lrr1* mutant, *kin7.lrr1* overexpressing KIN7-GFP, KIN7-DD-GFP or KIN7-AA-GFP *Arabidopsis* roots. Seedlings were grown for 6 days on MS plates. Both *kin7.lrr1* and *kin7.lrr1* expressing KIN7-AA present a defect in callose deposition with reduced levels internal tissues and in epidermal cells, compared to the Col-0. In the opposite way, overexpression of KIN7 and KIN7-DD phosphomimic induces an increase in callose deposition. (A) Representative confocal images of callose immunofluorescence (red) in roots. DAPI staining of DNA (blue) was performed to highlight the cellular organisation of root tissues. (B) Callose quantifications in “internal” root cell layers and epidermal cells. C, LR number in wild type Col-0 and PdBG1 overexpressing line. *Arabidopsis* lines were grown for 9 days on MS plates for control conditions, or 6 days then transferred to MS plate containing 0.4 M mannitol before root phenotyping. LR number is represented by white and red box plots for control and mannitol treatment, respectively. In control conditions, PdBG1 over expressor displays a decrease of LR number compared to the wild type. In response to mannitol treatment, Col-0 wild type and *Arabidopsis* seedlings overexpressing PdBG1 showed a decrease in LR number. The primary root length was measured in parallel to the LR (A) using FIJI software. None of the lines tested presented a significant root length difference compare to Col-0 in control conditions (white box plot). After mannitol treatment, all the lines were similarly affected with a reduction of the primary root length (red box plot).

Taking together, these results suggest that KIN7 is necessary to regulate LR development and response to mannitol *via* a mechanism possibly involving the synthesis and/or degradation of plasmodesmata-associated callose.

### KIN7 plasmodesmata localization is required to regulate callose and the root response to mannitol

Changes in KIN7 phosphorylation were found to be necessary for localisation of the protein at plasmodesmata in response to mannitol. To investigate the implications of KIN7 phosphorylation for LR response to mannitol, we tested complementation of *kin7.lrr1* phenotype with both the KIN7 phosphomimic (KIN7-DD) and the phosphodead (KIN7-AA) mutant variants. Under control conditions, over expression of both KIN7-DD-GFP and KIN7-AA-GFP variants in the *kin7.lrr1* mutant background did not affect root length (Fig. 6B, white box plots). Reduced LR phenotype in *kin7.lrr1* mutant was fully restored by expression of KIN7-DD, and only partially by expression of KIN7-AA (Fig. 6A, white boxes). Concomitantly, lines expressing KIN7-AA variant displayed a delay in LR primordium development with no stage VI and VII primordia at 42h after gravistimulation, a phenotype resembling *kin7.lrr1* (Fig. 6C). Next, we tested the phenotype of these lines in mannitol. As in wild type Col-0, LR number was reduced in response to mannitol in *kin7.lrr1* mutants expressing the phosphomimic but not the phosphodead KIN7 variant suggesting that KIN7 phosphorylation is important for LR response to mannitol (Fig.6A, compare white to red boxes).

We previously saw a defect in callose regulation at plasmodesmata in the *kin7.lrr1* (Fig.8), so we next investigated the effect of KIN7 phosphomimic and phosphodead variants on the callose mutant phenotype. We used immunolocalization to compare callose levels in wild type and in the *kin7.lrr1* mutant expressing either KIN7-AA or KIN7-DD (Fig.8A-B). While callose levels in the *kin7.lrr1* mutant expressing the phosphomimic version were comparable to KIN7 over expressing line, the phosphodead variant displayed a reduction of callose levels comparable to *kin7.lrr1* mutant (Fig. 8A-B).

To summarize, expression and phosphorylation-dependent relocalisation of KIN7 is important to regulate LR response to mannitol *via* a mechanism that modulates the levels of callose.

## DISCUSSION

In this study we report the rapid change of location of two PM-located LRR-RLKs in response to osmotic stress. Under standard growth conditions, both KIN7 and IMK2 show an exclusive PM localisation, but exposure to salt or mannitol triggered their relocalisation to plasmodesmata. This re-arrangement happens remarkably fast, within the first two 2 minutes after stimulation, suggesting that this process may be either post-transcriptionally or post-translationally regulated. Dynamic plasmodesmal association is neither a general feature of PM-associated proteins nor of microdomain-associated proteins, such as REM1.2 and 1.3, which localisations remain “static”. So far receptor-like proteins that associate with plasmodesmata have been reported to be spatially and stably confined to the PM microdomain lining the pores (Caillaud et al., 2014; Carella et al., 2015; Faulkner et al., 2013; Lim et al., 2016; Stahl et al., 2013a; Thomas et al., 2008; Vaddepalli et al., 2014). Conditional association with plasmodesmata have however been reported for the ER-PM membrane contacts site protein, Synaptotagmin SYTA, which within few days post-viral infection is recruited by *Tobamovirus* viral movement protein to plasmodesmata active in cell-to-cell spread (Levy et al., 2015). Our data reporting rapid re-organisation of two LRR-RLKs, suggests that plasmodesmata molecular composition is more dynamic than previously thought and most likely changes in response to environmental stimuli.

An important feature of the PM, which acts at the interface between the apoplastic and symplastic compartment, is its ability to respond to external and internal stimuli by remodelling its molecular organisation. This process takes many forms from the association/dissociation of proteins with nano-domains and complexes, through protein/protein and protein/lipid interactions, through the modification of ER-PM contacts, or post-translational modification such as phosphorylation or ubiquitination (Demir et al., 2013; Dubeaux et al., 2018; Julien Gronnier et al., 2017; Lee et al., 2019; Perraki et al., 2018). This, most likely also applies to plasmodesmata, which need to quickly integrate development and biotic/ abiotic stimuli to regulate their aperture. Spatio-temporal re-arrangement of RLKs from the bulk PM to plasmodesmata may provide a different membrane environment and protein partners, which in turn could modify the protein function. In line with that, the RLK CRINKLY4, is known to interact with CLAVATA1 and the heteromer displays different composition at the PM and at plasmodesmata indicating that local territory indeed modifies receptor activity/function (Stahl et al., 2013).

In plants, protein mobility within the plane of the PM is restricted by the cell wall and appears to be rather slow compared to animal cells (Martiniere et al., 2012). Rapid re-arrangement of KIN7 within the plane of the PM was therefore unexpected. This pushed us to investigate the molecular determinants controlling plasmodesmata association. Our group previously showed that the specialised PM domain of plasmodesmata is enriched in sterols and sphingolipids. Altering the membrane sterol pool lead to plasmodesmata protein mis-localisation and defcets in callose-mediated cell-to-cell trafficking (Grison et al. 2015a). Both KIN7 and IMK2 were reported to associate with DRM (Demir et al., 2013; Keinath et al., 2010; Kierszniowska S, Seiwert B, 2009; Shahollari et al., 2005; Srivastava et al., 2013; Szymanski et al., 2015), hence supposedly sterol- and sphingolipid-enriched PM nanodomains. However, inhibiting sterol- and VLCFA-sphingolipid synthesis had no effect on KIN7 relocalisation to plasmodesmata upon stress conditions (Demir et al., 2013; Kierszniowska S, Seiwert B, 2009).

Protein phosphorylation has been reported as one of the early post-translational responses to osmotic stress (Nikonorova et al., 2018) and KIN7 has multiple phosphorylation sites and is phosphorylated in response to abiotic stress (Chang et al., 2012; Niittylä et al., 2007). Using phospho-mutants of KIN7, we showed that the phosphorylation status of KIN7 is important for subcellular localisation with the KIN7-DD phosphomimic mutant partially associating with plasmodesmata even in control conditions, while the KIN7-AA phosphodead mutant was significantly affected in its capacity to localise to plasmodesmata after mannitol treatment. Having said that, KIN7-AA mutant is still able to partially localise to the pores after stress (PD index of 1.2) indicating that other factors may be important to control this process. For KIN7, localization to the PM microdomains was previously shown to depend on cytoskeletal integrity (Szymanski et al., 2015) and involvement of cytoskeletal components in re-organisation to plasmodesmata should be investigated in further studies.

An explanation for why KIN7 and IMK2 cluster at plasmodesmata in response to mannitol and NaCl, and how this exactly impact on plasmodesmata function remains to be determined. We postulate that our mannitol treatment induces a change in plasmodesmata permeability through callose deposition or removal as it has been observed for cold, oxidative, nutrient, and biotic stresses (Benitez-Alfonso et al., 2011; Bilska and Sowinski, 2010; Cui and Lee, 2016; Faulkner et al., 2013; Lexy et al., 2018; Sivaguru et al., 2000; Zavaliev et al., 2011). Callose is a well-established regulator of plasmodesmata-mediated cell-to-cell communication and modifying callose deposition at the pores has a strong impact on numerous developmental programs including LR formation (Benitez-Alfonso et al., 2013; Maule et al., 2013; Otero et al., 2016). The balance between callose synthesis and degradation is tightly regulated through a set of callose-related enzymes. The plasmodesmata associated β1-3 glucanase PdBG1 (AT3G13560) is involved in modulating plasmodesmata aperture through callose degradation and has been implicated in LR formation and patterning (Benitez-Alfonso et al., 2013; Maule et al., 2013). Our data indicate that the KIN7 induced LR response in control and mannitol stress condition is likely to involve callose. Modifying plasmodesmata permeability by over-expressing PdBG1 affect LR phenotype and resembles that of *kin7.lrr1* and *kin7.lrr1* over-expressing KIN7-AA lines, which are also defective in callose regulation.

To conclude, our work highlights the complex and dynamic regulation of symplastic intercellular communication in response to osmotic stress, a situation that plants are often confronted to in their environment. We propose that re-organisation of PM-located RLKs to plasmodesmata is an ingenious mechanism which combines “stress sensing” at the bulk PM and modulation of cell-to-cell trafficking at plasmodesmata.

## SUPPLEMENTAL FIG.S

**Supplemental Figure 1.**
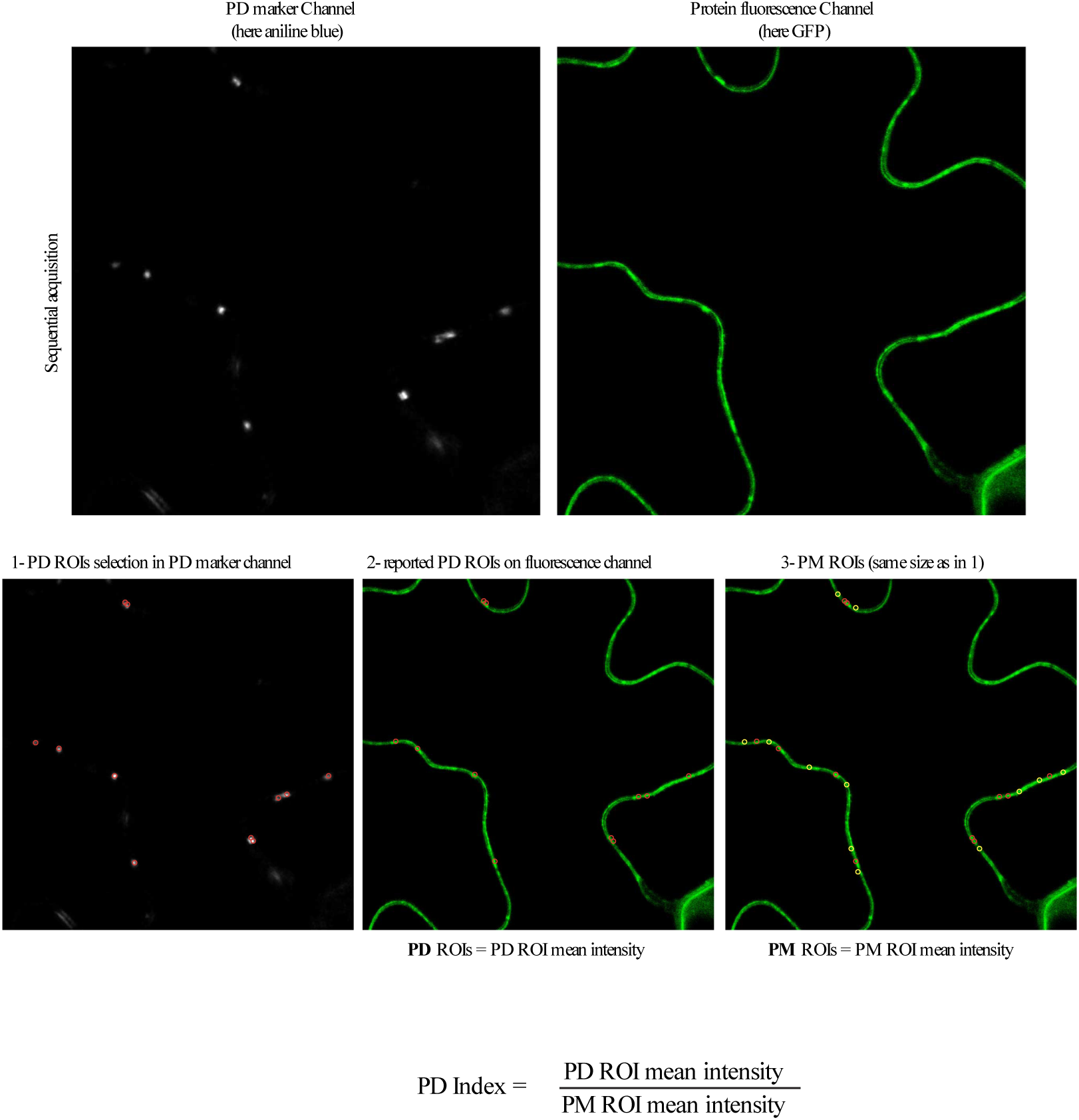
PD Index calculation. Plasmodesmata depletion or enrichment was assessed by calculating for a given protein the fluorescence intensity ratio between plasmodesmata (indicated PDLP1-mRFP or aniline blue; red circles/ROIs) versus the plasma membrane outside plasmodesmata (yellow circles/ROIs). A PD index above 1 indicate plasmodesmata enrichment. PD, plasmodesmata; PM, plasma membrane; ROI, region of interest.

**Supplemental Figure 2.**
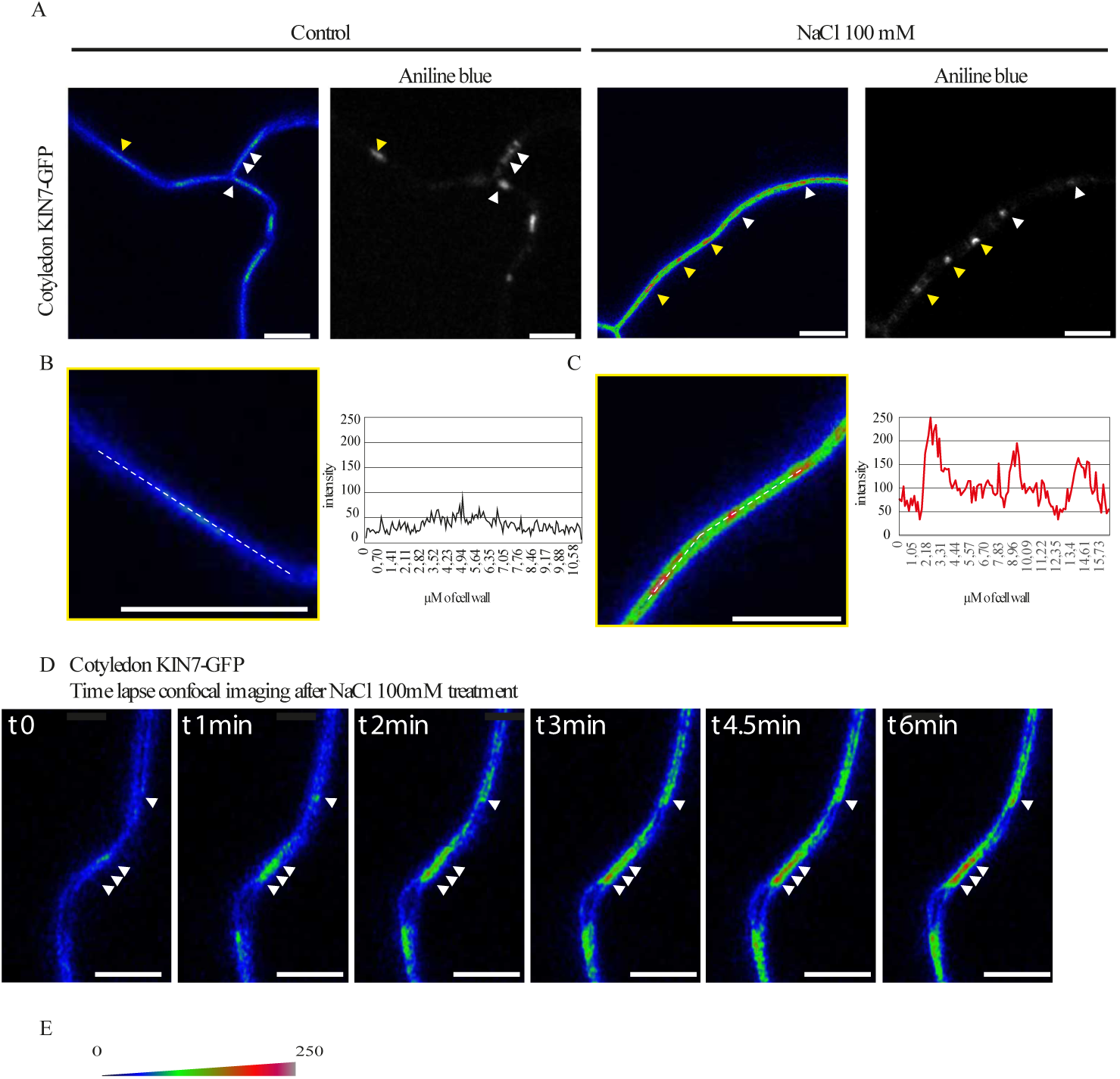
Stable *Arabidopsis* line expressing KIN7-GFP, under 35S promoter and visualised by confocal microscopy. All images have been color-coded through a heat-map filter to highlight clustering at plasmodesmata. A-D, In control conditions, KIN7-GFP localises exclusively at the PM in cotyledons (A-C) and is not enriched at plasmodesmata (marked by aniline blue staining, arrowheads). B and C are magnified regions indicated by yellow arrowheads in A. Upon NaCl 100 mM (1-5 min), KIN7 relocalises to plasmodesmata where it becomes enriched (A, arrowheads). Intensity plots along the white dashed lines are shown for KIN7-GFP localisation pattern in control and NaCl conditions. D, Time-lapse imaging of KIN7-GFP relocalisation upon NaCl exposure. Within less than two minutes plasmodesmata localisation already visible (white arrowhead). E, Shows a color-coding bar for heat-map images. Scale bars= 10 μm

**Supplemental Figure 3.**
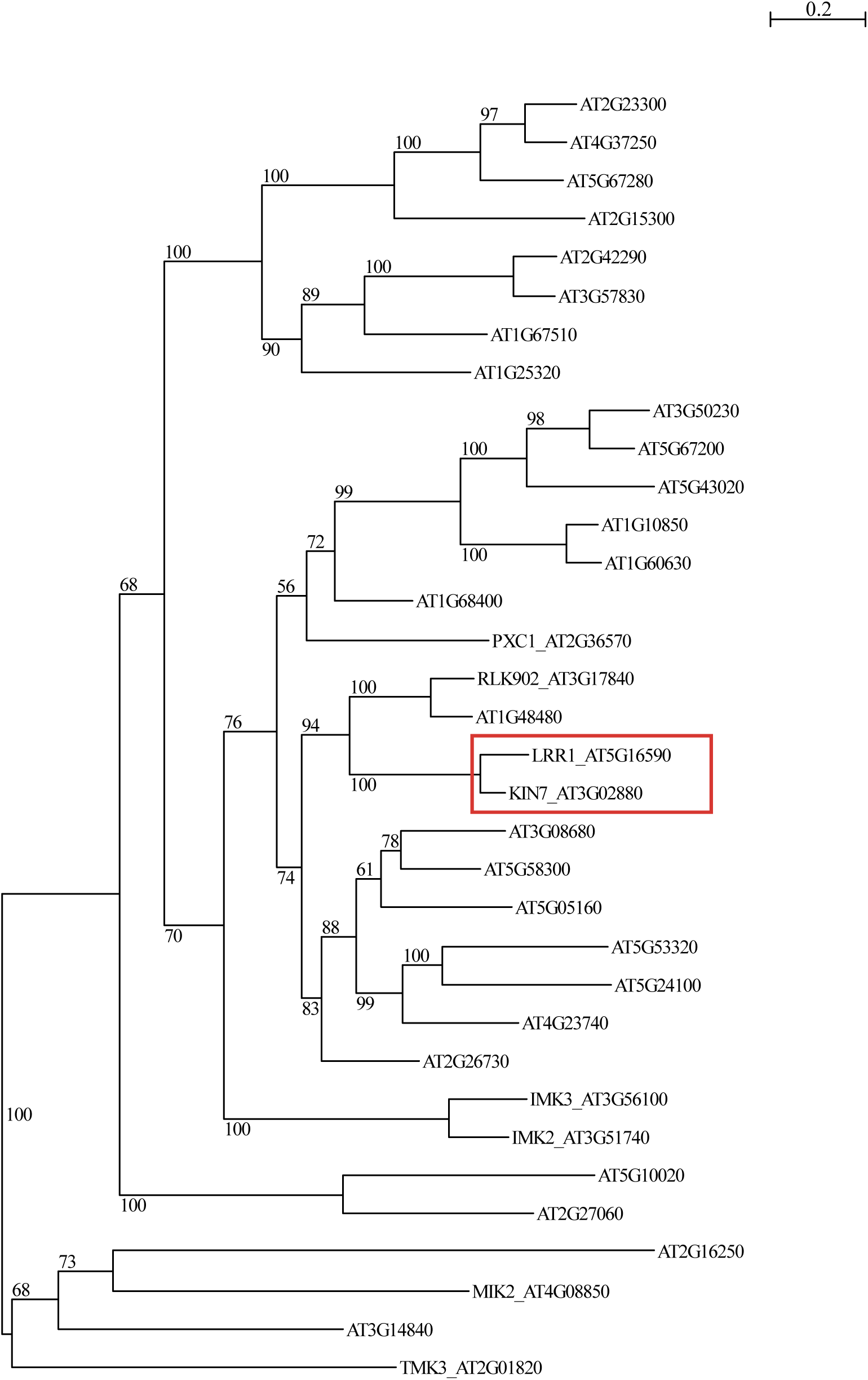
Clade III LRR-RLK subclade PhyML ln(L)=-14326.2 320 sites LG 1000 replic. 4 rate classes. Phylogenic tree of clade III LRR-RLKs showing that KIN7 and LRR1 are closely related.

**Supplemental Figure 4.**
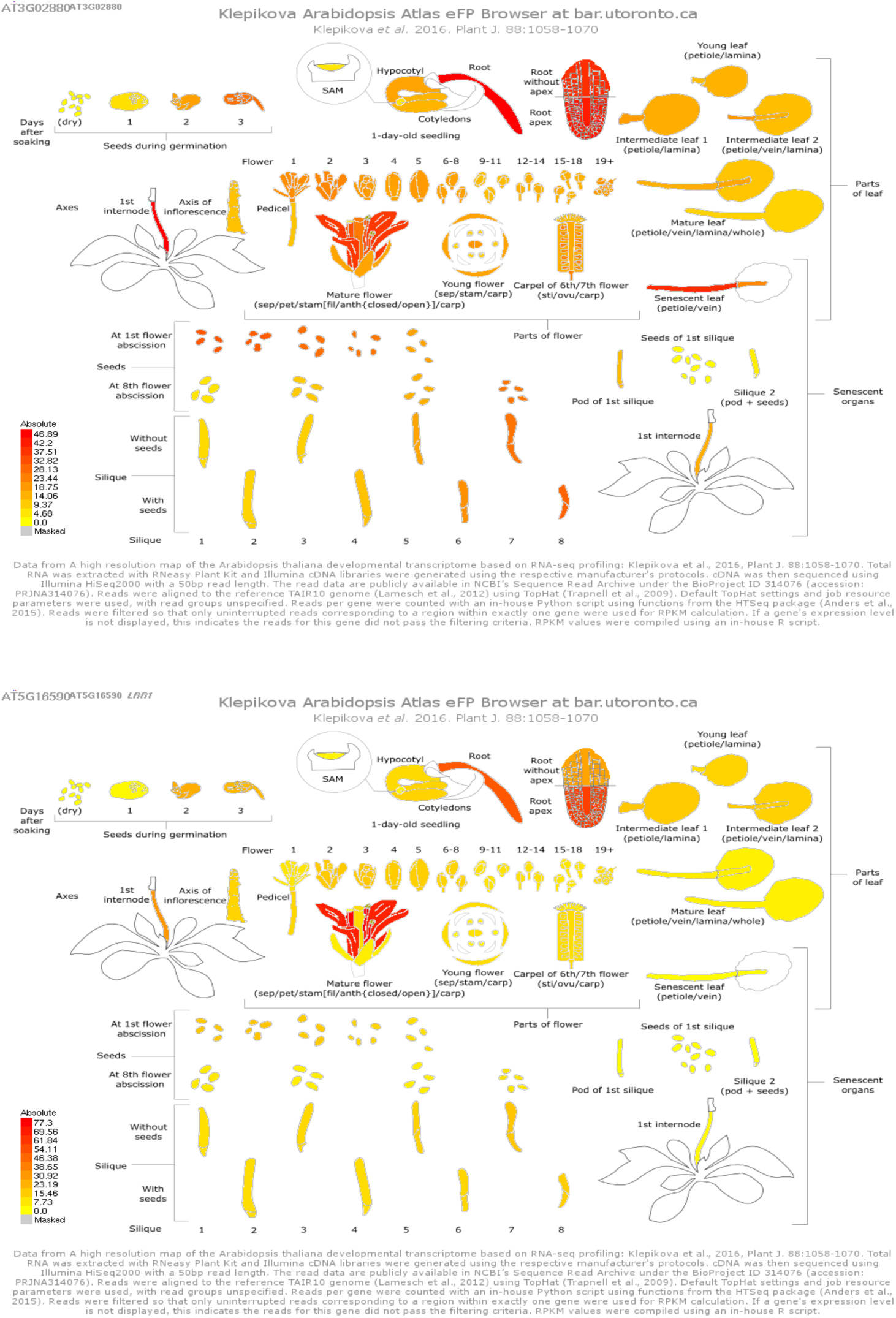
KIN7 and LRR1 display similar expression pattern. Profile extrated from the Bio-Analytic Ressource for Plant Biology. Expression pattern of KIN7 and LRR1 extracted from the Bio-Analytic Ressource for Plant Biology (bar.utoronto.ca) based on developmental transcriptome based RNA-seq profiling (Klepikova et al., 2016) showing similar expression patterns.

**Supplemental Figure 5.**
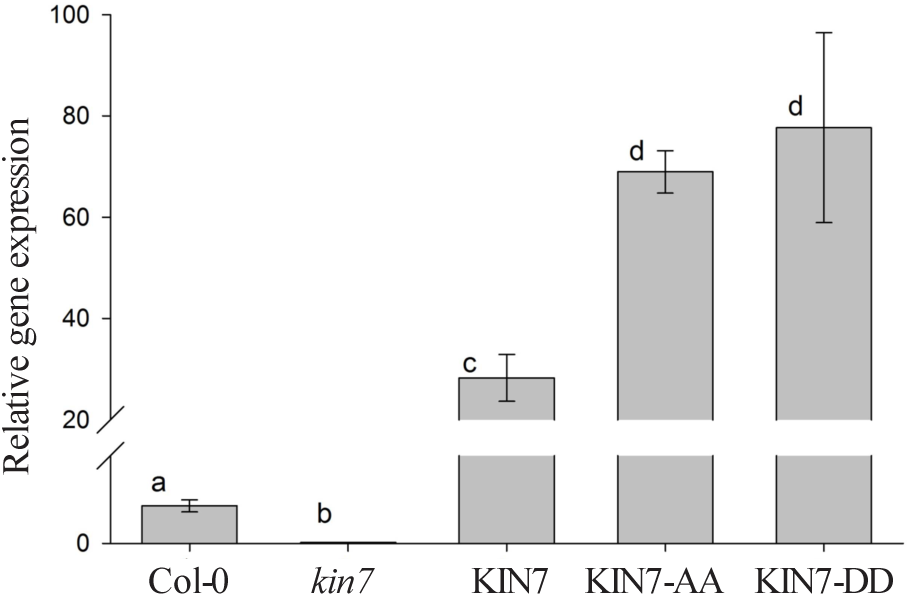
Expression of KIN7, and KIN7-GFP, KIN7-DD-GFP and KIN7-AA-GFP transgenes in *kin7.lrr1* mutant background.

**Supplemental movie 1**

Time lapse confocal movie showing the rapid re-localisation of KIN7-GFP immediately after mannitol treatment. Time scale is visible at the top left. Color-coding bar for heat-map images same as in Figure 2.

**Supplemental table S1.**
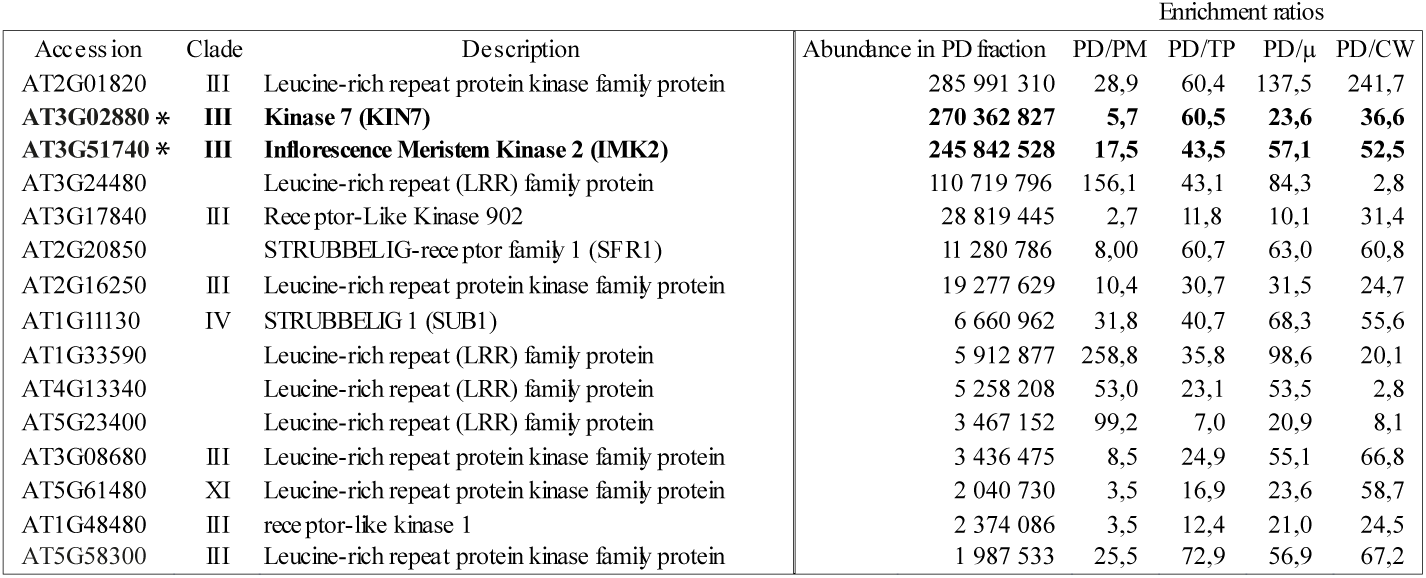
List of RLKs extracted from the label-free *Arabidopsis* plasmodesmata proteome from Brault et al., 2018. PD, plasmodesmata fraction; TP; total cellular protein fractions, μ, microsomal protein fraction; CW, cell wall protein extracts. Stars: LRR-RLKs selected for further localisation analysis.

**Supplemental table S2.**
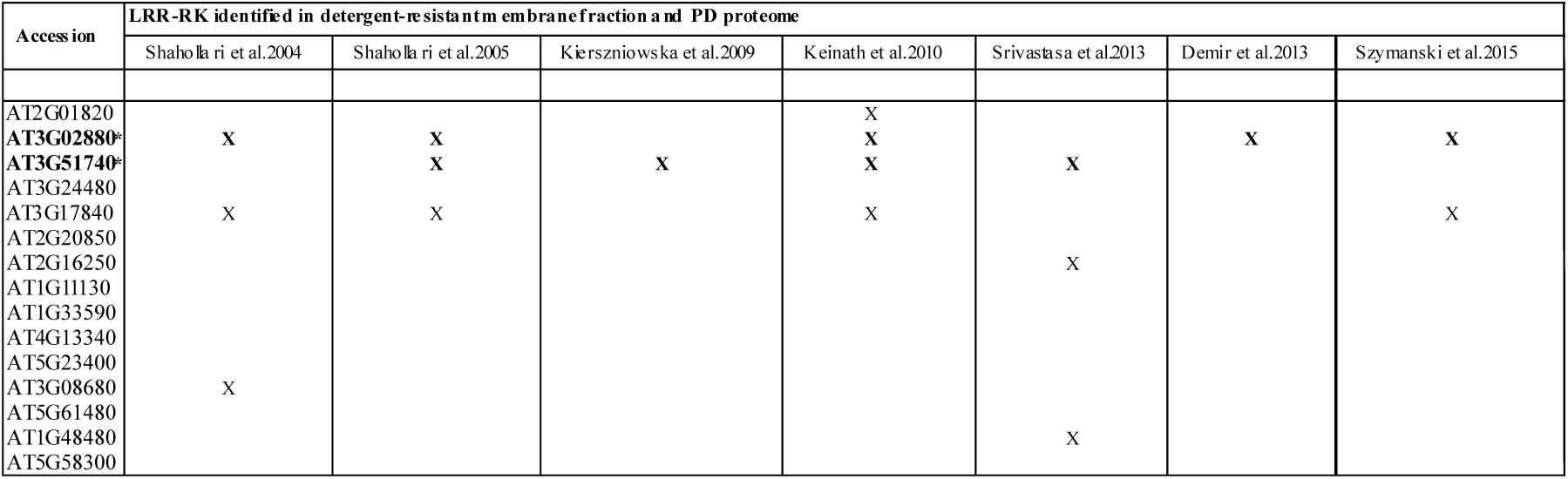
RLKs associated with lipid microdomains according to seven Detergent Resistant Membrane proteomic studies. The list of RLKs present in the *Arabidopsis* plasmodesmata proteome (Supplementary Table S1) was crossed referenced with published Detergent Resistant Membrane proteomes. RLKs were selected when present in at least two independent proteomic studies.

**Supplemental Table S3.**
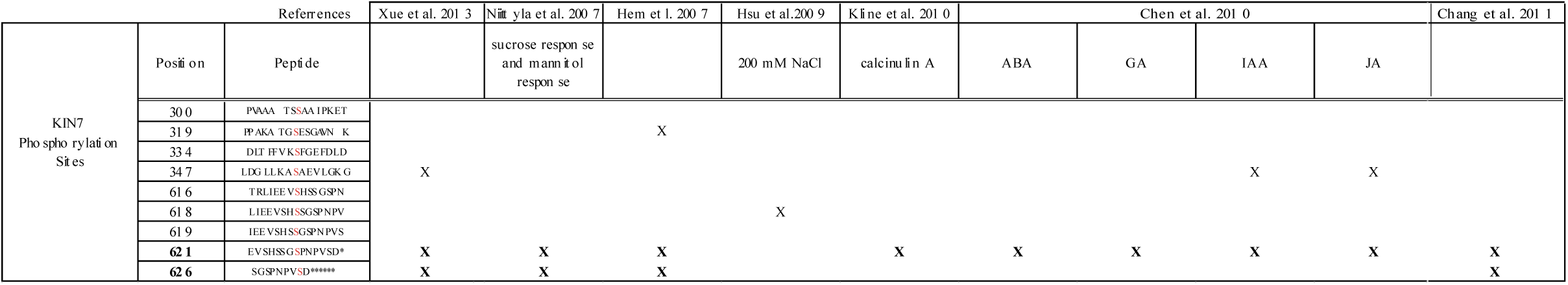
KIN7 phosphorylation sites (indicated in red) detected in phosphoproteomic studies. In bold the two phosphor-sites selected for this study. Stars indicate the end of the protein.

**Supplemental table 4.**
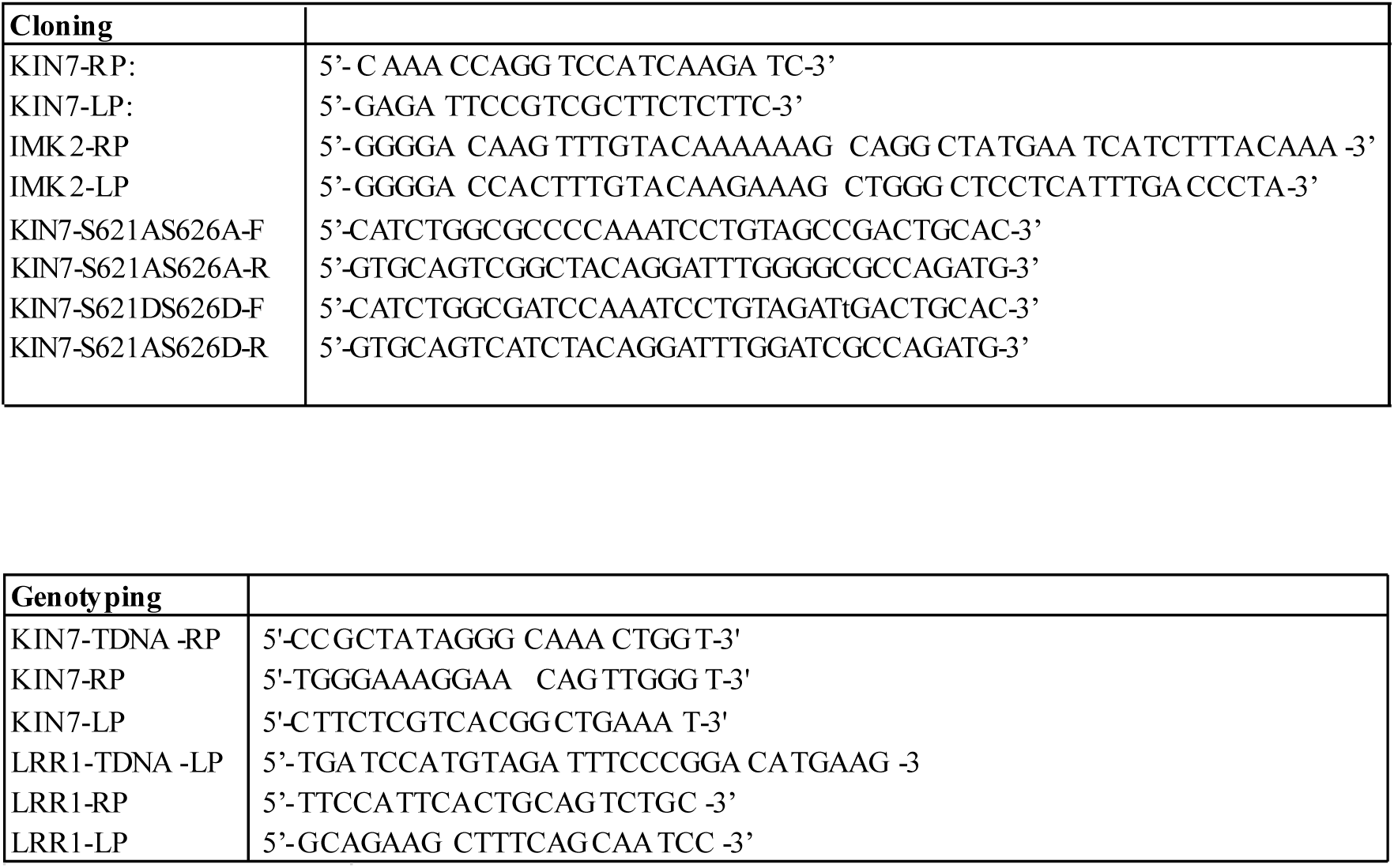
Primer list. List of primers used in the present work

## Acknowledgements

This work was supported by the National Agency for Research (Grant ANR-14-CE19-0006-01 to E.M.B), “Osez l’interdisciplinarité” OSEZ-2017-BBRIDGING CNRS program to E.M.B., the European Research Council (ERC) under the European Union’s Horizon 2020 research and innovation programme (grant agreement No 772103-BRIDGING to E.M.B). P.K. was supported by a BBSRC DTP (BB/M011151/1). Y.B.-A. lab work is supported by research grants from the Leverhulme Trust RPG-2016-136. Work in W.X.S. lab was funded by the Deutsche Forschungsgemeinschaft, grant SCHU1533/9-1 to WS and XW.

Fluorescence microscopy analyses were performed at the plant pole of the Bordeaux Imaging Centre (http://www.bic.u-bordeaux.fr). The lipidomic analyses were performed at the Functional Genomic Center of Bordeaux, Metabolome/Lipidome platform (https://metabolome.cgfb.u-bordeaux.fr/en) funded by Grant MetaboHUB-ANR-11-INBS-0010.

We thank Jens Tilsner for critical review of the article prior to submission.

## Contributions

M.S.G. performed all experiments and analysed data, with the exception of *kin7.lrr1* mutant and KIN7-GFP, KIN7-AA and KIN7-DD transgenic Arabidopsis lines, which were generated by X.N.W. M.L.B. helped with NaCl image acquisition and callose quantification. F.I. helped with proteomic analysis and cross-references with published proteomic data sets and phylogenetic tree. Y.B.A and P.K. made a substantial contribution to carrying out the study by performing research described in Fig. 7D-E and Fig. 8C. Y.B.A. also contributed to the analysis and interpretation of study data, helped draft the output and critique the output for important intellectual content.

E.M.B. and M.S.G. designed the research with the help of F.I and Y.B.A.. E.M.B and M.S.G. wrote the manuscript with the help of of F.I and Y.B.A. All the authors discussed the results and commented on the manuscript.

## Material and methods

### Proteomic analyses

We used the label-free plasmodesmata proteomic analysis of Brault et al. (Brault et al., 2018) to select RLK candidates. For that all members of the LRR-RLK family which displayed with a significant fold change (plasmodesmata/PM enrichment ratio >2) were selected (Supplemental Table. S1) and crossed reference with DRM proteomic studies (Supplemental Table. S2).

### Cloning

IMK2 and KIN7 were cloned using classical gateway system with p221 as DNR plasmid and pGBW661 or pGBW641 as DEST plasmid comprising 35S promoter and C terminal tag GFP and TagRFP respectively. KIN7-AA and KIN7-DD were cloned using primers in supplemental table S4). Amplifications were run on plasmid containing the full-length cDNA (U12366 TAIR), purified with QIAquick gel extraction kit and inserted into p221 DNR (See Supplemental Table. S4 for primer details) and then inserted into pDEST for stable expression in *A*. *thaliana* or for transient expression in *N. benthamiana*.

### Plant Material and Growth Conditions

The following *Arabidopsis* transgenic lines were used: p35S:Lti6b-mCherry; p35S::PIP2;1-GFP; pREM1.2:REM1.2-YFP, pREM1.3:REM1.3-YFP, p35S::PdBG1 (Benitez-Alfonso et al., 2013; Cutler et al., 2000; Jarsch et al., 2014; Prak et al., 2008; Szymanski et al., 2015).

Generation of *kin7*.*lrr1* loss-of-function Arabidopsis mutants and overexpressing KIN7 lines: *Kin7* (SALK_019840) and *lrr1* (WiscDsLoxHs082_03E) T-DNA insertional Arabidopsis mutants (background Col-0) were obtained from the Arabidopsis Biological Resource Center (http://www.arabidopsis.org/). Single T-DNA insertion lines were genotyped and homozygous lines were crossed to obtain double homozygous *kin7.lrr1*.

T-DNA insertional mutants *kin7, lrr1* and double mutant *kin7.lrr1* were confirmed via PCR amplification using T-DNA border primer and gene specific primers (Supplemental Table S4). For genotyping, genomic DNA was extracted from Col-0, *kin7.lrr1* plants using chloroform:isoamyl alcohol (ratio24:1), genomic DNA isolation buffer (200mM Tris HCL PH7.5, 250mM NaCl, 25mM EDTA and 0.5% SDS) and isopropanol. PCR were performed with primers indicated in Supplemental Table. S4.

We generated p35S:KIN7-GFP, p35S:KIN7-S621D_S626D-GFP and p35S:KIN7-S621A_S626A-GFP in *kin7.lrr1* mutant background. Lack of KIN7 expression in the double mutant background and overexpression of KIN7-GFP KIN7-DD and KIN7AA was demonstrated by RT-PCR (Supplementary Fig. S5). For that, total mRNA was extracted from Arabidopsis line using RNeasy® Plant Mini Kit (QIAGEN) and cDNA was produced using random and oligodT primers.

For confocal microscopy, *Arabidopsis* seedlings were grown 6 days on agar plate 8g/L containing MS salts including vitamins 2,2g/L, sucrose 10g/L and MES 0,5g/L at pH 5,8 in a culture room at 22°C in long day light conditions (150µE/m^2^/s) followed by treatment with NaCl or mannitol (see below for details).

For LR phenotyping, *Arabidopsis* seedlings were grown 9 days on agar plate 8g/L containing MS salts including vitamins 2,2g/L, sucrose 10g/L and MES 0,5g/L at pH 5,8 in a culture room at 22°C in long day light conditions (150µE/m^2^/s) for control conditions or 6 days then transferred to the same media supplemented with mannitol 0.4M for another 3 days.

### Mannitol and NaCl treatments

For short-term treatment, mannitol (0.4 M solution) or NaCl (100 mM solution) were infiltrated in *Arabidopsis* cotyledons (for stable expression) or *N. benthamiana* leaves (for transient expression), and samples were immediately observed by confocal microscopy. For *Arabidopsis* roots, seedling were grown for 6 days on ½ MS 1% sucrose agar plates in long day conditions then transferred in liquid ½ MS 1% sucrose media containing 0.4 M mannitol for 3h before analysis (confocal live imaging or immunolocalisation against calloseon whole mount tissues). For control conditions, leaves/cotyledons were infiltrated with water and *Arabidopsis* roots incubated in ½ MS 1% sucrose media without mannitol. For long-term mannitol treatment, seedlings were grown for 6 days on ½ MS 1% sucrose agar plates in long day conditions, then transferred on ½ MS 1% sucrose agar plates containing 0.4M of mannitol for 3 days, before analysis

### Confocal live imaging

For transient expression in *N. Benthamiana*, leaves of 3 week-old plants were pressure-infiltrated with GV3101 agrobacterium strains, previously electroporated with the relevant binary plasmids. Prior to infiltration, agrobacteria cultures were grown in Luria and Bertani medium with appropriate antibiotics at 28°C for one days then diluted to 1/10 and grown until the culture reached an OD_600_ of about 0.8. Bacteria were then pelleted and resuspended in water at a final OD_600_ of 0.3 for individual constructs, 0.2 each for the combination of two. Agroinfiltrated *N. benthamiana* leaves were imaged 3 days post infiltration at room temperature using a confocal laser scaning microscope Zeiss LSM 880 using X63 oil lens. Immediately before imaging leaves were infiltrated with H_2_O, 0.4 M mannitol or 100 mM NaCl solutions supplemented with 20 µg/mL aniline blue (Biosupplies) for plasmodesmata co-localisation and PD index, ∼ 0.5cm leaf pieces were cut out and mounted with the lower epidermis facing up onto glass microscope slides.

For *Arabidopsis* lines, seedlings were grown for 6 days on ½ MS 1% sucrose agar plate prior to treatment. For cotyledon observation, seedlings were vacuum infiltrated with H_2_O or 0.4 M mannitol treatment supplemented with 20 µg/mL aniline blue and immediately mounted onto glass microscope slides with the lower epidermis facing up for confocal observation. For roots, seedling were incubated for 3h with appropriate solution before observation.

For time-lapse imaging, KIN7 expressing *Arabidopsis* cotyledons were cut in half and dry mounted onto microscope glass and cover slip, and 0.4 M mannitol solution was gently injected between glass and cover slip, and immediately followed by imaging.

For GFP and YFP imaging, excitation was performed with 2-8% of 488 nm laser power and fluorescence emission collected at 505-550 nm and 520-580 nm, respectively. For mRFP imaging, excitation was achieved with 2-5% of 561 nm laser power and fluorescence emission collected at 580-630 nm. For aniline blue imaging, excitation was performed with 0,5 to 6% of 405 nm laser power and fluorescence emission collected at 420-480 nm. For co-localisation sequential scanning was systematically used.

### PD index

Plasmodesmata depletion or enrichment was assessed by calculating the fluorescence intensity ratio between the GFP/YFP/mRFP/mCherry-tagged protein intensity at plasmodesmata (indicated PDLP1-mRFP or aniline blue) versus the plasma membrane outside plasmodesmata. Confocal images of leaf/cotyledon or roots epidermal cells (*N. benthamiana* or *Arabidopsis*) were acquired by sequential scanning of PDLP1-mRFP or aniline blue (as plasmodesmata markers) and GFP/YFP/mRFP/mCherry-tagged (for confocal setting see above). About thirty images of leaf epidermis cells were acquired with a minimum of three biological replicates. Individual images were then processed using Fiji by defining five to twenty regions of interest (ROI) at plasmodesmata (using plasmodesmata marker to define the ROI) and five to twenty ROIs outside plasmodesmata. The ROI size and imaging condition were kept the same. The GFP/YFP/mRFP/mCherry-tagged protein mean intensity was measured for each ROI then averaged for single image. The plasmodesmata index corresponds to intensity ratio between fluorescence intensity of proteins at plasmodesmata versus outside the pores. (see Supplemantal Fig. S1)

### Callose quantification in Arabidopsis roots by whole-mount immunolocalisation

Arabidopsis seedlings were grown on ½ MS 1% sucrose agar plate for 6 days then incubated 3 hours in ½ MS 1% sucrose liquid media for control condition or ½ MS 1% sucrose liquid media containing 0.4 M mannitol, prior to fixation. The immunolocalization procedure was done according to Boutté *et al*. 2014 (Boutté and Grebe, 2014). The callose antibody (Australia Biosupplies) was diluted to 1/300 in MTSB (Microtubule Stabilizing Buffer) containing 5% of neutral donkey serum. The secondary anti-mouse antibody coupled to TRITC (tetramethylrhodamine) was diluted to 1/300 in MTSB buffer containing 5% of neutral donkey serum. The nucleus were stained using DAPI (4’,6-diamidino-2-phénylindole) diluted to 1/200 in MTSB buffer for 20 minutes. Samples were then imaged with a Zeiss LSM 880 using X40 oil lens. DAPI excitation was performed using 0,5% of 405 laser power and fluorescence collected at 420-480 nm; GFP excitation was performed using 5% of 488 nm laser power and fluorescence emission collected at 505-550 nm; TRITC excitation was performed with 5% of 561 nm power and fluorescence collected at 569-590 nm. All the parameters were kept between experiments to allow quantifications.

Callose deposition was then quantified using Fiji software. Callose fluorescence intensity was measured at the apico-basal cell walls of epidermal cells and internal layers endodermal and cortex cells for the “inner tissues”. A total of 20 cell wall intensity were measured per cell lineage (e.g. 20 epidermal; 20 endodermal + 20 cortex) per roots, 10 roots per transgenic lines. Two biological replicate were done.

### LR number and LR primordium developmental stage quantifications

Arabidopsis seedling were grown 9 days on ½ MS 1%sucrose agar plates for control or 6 days on ½ MS 1% sucrose agar plates then transferred for 3 days on ½ MS 1% sucrose agar plates supplemented with 0.4 M mannitol. The number of emerged LRs and LR primordia (from stage 2) was imaged and quantified using a macroscope Axiozoom Leica with a 150X magnification. LR primordium stages were analysed according to (Péret et al., 2012).

Root length was measured by using Image J software after taking pictures of the plates with Biorad Chemidoc.

### Sterol and sphingolipid inhibitor Treatments

For sterols and sphingolipids inhibitor experiments, 5 days-old seedlings were transferred to MS agar plates containing 100 μg/mL Fenpropimorph (stock solution 100 mg/mL in DMSO) or 100 nM Metazachlor (stock solution 1 mM in DMSO). Control plates contained an equal amount of 0.1% DMSO solvent. Seedlings were observed by confocal microscopy 48h after treatment and lipid analysis was performed in parallel (see below for details).

### Lipid Analysis

For the analysis of total fatty acids by GC-MS (FAMES), Arabidopsis seedlings were harvested 48h after transfer on MS plates containing 100nM Metazachlor or 0.1%DMSO. Transmethylation and trimethylsilylation of fatty acids from 150mg of fresh material was performed as describe in (Magali S. Grison et al., 2015). An HP-5MS capillary column (5%phenyl-methyl-siloxane, 30-m, 250-mm, and 0.25-mm film thickness; Agilent) was used with helium carrier gas at 2 mL/min; injection was done in splitless mode; injector and mass spectrometry detector temperatures were set to 250°C; the oven temperature was held at 50°C for 1 min, then programmed with a 25°C/min ramp to 150°C (2-min hold) and a 10°C/min ramp to 320°C (6-min hold). Quantification of non-hydroxylated and hydroxylated fatty acids was based on peak areas that were derived from the total ion current.

For sterols analysis by GC-MS, Arabidopsis seedlings were harvested 48h after transfer on MS plates containing 100µg/mL Fenpropimorph or 0.1%DMSO. A saponification of 150mg of fresh material was performed by adding 1 mL of ethanol containing the internal standard α-cholestanol (25µg/mL) and 100 mL of 11 N KOH and incubating it for 1 h at 80°C. After the addition of 1 mL of hexane and 2 mL of water, the sterol-containing upper phase was recovered and evaporated under an N2 gas stream. Sterols were derivatized by BSTFA as described for FAMEs and resuspended in 100 µL of hexane before analysis by GC-MS (see FAME analysis).

### Phylogenetic Tree Construction

Sequence alignment and phylogenetic tree building were performed with SeaView version 4 multiplatform program. Alignment algorithm chosen was ClustalW and PhyML version 3 was used to reconstruct maximum-likelihood tree of 34 clade III LRR-RLKs (Hove et al., 2011)

### Statistical analysis

Statistical analyses were done using “R” software. For all analyses, we applied “Wilcoxon rank sum test” which is a non-parametrical statistical test commonly used for small range number of replicate (e.g. n<20).

## REFERENCES

Amari, K., Boutant, E., Hofmann, C., Schmitt-Keichinger, C., Fernandez-Calvino, L., Didier, P., Lerich, A., Mutterer, J., Thomas, C.L., Heinlein, M., Mély, Y., Maule, A.J., Ritzenthaler, C., 2010. A family of plasmodesmal proteins with receptor-like properties for plant viral movement proteins. PLoS Pathog. 6, 1–10. https://doi.org/10.1371/journal.ppat.1001119

Bayer, E., Thomas, C.L., Maule, a J., 2004. Plasmodesmata in Arabidopsis thaliana suspension cells. Protoplasma 223, 93–102. https://doi.org/10.1007/s00709-004-0044-8

Benitez-Alfonso, Y., Faulkner, C., Pendle, A., Miyashima, S., Helariutta, Y., Maule, A., 2013. Symplastic Intercellular Connectivity Regulates Lateral Root Patterning. Dev. Cell 26, 136–147. https://doi.org/10.1016/j.devcel.2013.06.010

Benitez-Alfonso, Y., Faulkner, C., Ritzenthaler, C., Maule, A.J., 2010. Plasmodesmata: gateways to local and systemic virus infection. Mol. Plant. Microbe. Interact. 23, 1403– 1412. https://doi.org/10.1094/MPMI-05-10-0116

Benitez-Alfonso, Y., Jackson, D., Maule, A., 2011. Redox regulation of intercellular transport. Protoplasma 248, 131–140. https://doi.org/10.1007/s00709-010-0243-4

Bilska, A., Sowinski, P., 2010. Closure of plasmodesmata in maize (Zea mays) at low temperature: a new mechanism for inhibition of photosynthesis. Ann. Bot. 106, 675–686. https://doi.org/10.1093/aob/mcq169

Bocharov, E. V., Lesovoy, D.M., Pavlov, K. V., Pustovalova, Y.E., Bocharova, O. V., Arseniev, A.S., 2016. Alternative packing of EGFR transmembrane domain suggests that protein-lipid interactions underlie signal conduction across membrane. Biochim. Biophys. Acta - Biomembr. 1858, 1254–1261. https://doi.org/10.1016/j.bbamem.2016.02.023

Boutté, Y., Grebe, M., 2014. Immunocytochemical fluorescent in situ visualization of proteins in arabidopsis. Methods Mol. Biol. 1062, 453–472. https://doi.org/10.1007/978-1-62703-580-4_24

Brault, M., Petit, J.D., Immel, F., Nicolas, W.J., Brocard, L., Gaston, A., Fouché, M., Hawkins, T.J., Crowet, J.-M., Grison, S.M., Kraner, M., Alva, V., Claverol, S., Deleu, M., Lins, L., Tilsner, J., Bayer, E.M., 2018. Multiple C2 domains and Transmembrane region Proteins (MCTPs) tether membranes at plasmodesmata. BioRxiv doi.org/10.1101/423905.

Bücherl, C.A., Jarsch, I.K., Schudoma, C., Robatzek, S., Maclean, D., Ott, T., Zipfel, C., Genome, P., National, S., Biology, C., 2017. Plant immune and growth receptors share common signalling components but localise to distinct plasma membrane nanodomains 1–28. https://doi.org/10.7554/eLife.25114

Cacas, J.-L., Buré, C., Grosjean, K., Gerbeau-Pissot, P., Lherminier, J., Rombouts, Y., Maes, E., Bossard, C., Gronnier, J., Furt, F., Fouillen, L., Germain, V., Bayer, E., Cluzet, S., Robert, F., Schmitter, J.-M., Deleu, M., Lins, L., Simon-Plas, F., Mongrand, S., 2016. Revisiting plant plasma membrane lipids in tobacco: A focus on sphingolipids. Plant Physiol. 170. https://doi.org/10.1104/pp.15.00564

Caillaud, M.C., Wirthmueller, L., Sklenar, J., Findlay, K., Piquerez, S.J.M., Jones, A.M.E., Robatzek, S., Jones, J.D.G., Faulkner, C., 2014. The Plasmodesmal Protein PDLP1 Localises to Haustoria-Associated Membranes during Downy Mildew Infection and Regulates Callose Deposition. PLoS Pathog. 10, 1–13. https://doi.org/10.1371/journal.ppat.1004496

Carella, P., Isaacs, M., Cameron, R.K., 2015. Plasmodesmata-located protein overexpression negatively impacts the manifestation of systemic acquired resistance and the longdistance movement of Defective in Induced Resistance1 in Arabidopsis. Plant Biol. 17, 395–401. https://doi.org/10.1111/plb.12234

Chang, I., Hsu, J., Hsu, P., Sheng, W., Lai, S., Lee, C., 2012. Comparative phosphoproteomic analysis of microsomal fractions of Arabidopsis thaliana and Oryza sativa subjected to high salinity. Plant Sci. 185–186, 131–142. https://doi.org/10.1016/j.plantsci.2011.09.009

Chen, Y., Hoehenwarter, W., Weckwerth, W., 2010. Comparative analysis of phytohormone-responsive phosphoproteins in Arabidopsis thaliana using TiO 2 -phosphopeptide enrichment and mass accuracy precursor alignment. Plant J. 63, 1–17. https://doi.org/10.1111/j.1365-313X.2010.04218.x

Corbesier, L., 2009. FT Protein Movement Contributes to 1030. https://doi.org/10.1126/science.1141752

Cui, W., Lee, J.-Y., 2016. Arabidopsis callose synthases CalS1/8 regulate plasmodesmal permeability during stress. Nat. Plants 2, 16034. https://doi.org/10.1038/nplants.2016.34

Cutler, S.R., Ehrhardt, D.W., Griffitts, J.S., Somerville, C.R., 2000. Random GFP::cDNA fusions enable visualization of subcellular structures in cells of Arabidopsis at a high frequency. Proc. Natl. Acad. Sci. 97, 3718–3723. https://doi.org/10.1073/pnas.97.7.3718

Daum, G., Medzihradszky, A., Suzaki, T., Lohmann, J.U., 2014. A mechanistic framework for noncell autonomous stem cell induction in Arabidopsis. Proc. Natl. Acad. Sci. U. S. A. 111, 14619–24. https://doi.org/10.1073/pnas.1406446111

Deak, K.I., Malamy, J., Genetics, M., 2005. Osmotic regulation of root system architecture. Plant J. 43, 17–28. https://doi.org/10.1111/j.1365-313X.2005.02425.x

Demir, F., Horntrich, C., Blachutzik, J.O., Scherzer, S., Reinders, Y., Kierszniowska, S., Schulze, W.X., Harms, G.S., Hedrich, R., Geiger, D., Kreuzer, I., 2013. Arabidopsis nanodomain-delimited ABA signaling pathway regulates the anion channel SLAH3. Proc. Natl. Acad. Sci. 110, 8296–8301. https://doi.org/10.1073/pnas.1211667110

Dubeaux, G., Neveu, J., Zelazny, E., Vert, G., 2018. Metal Sensing by the IRT1 transporter-receptor orchestrates its own degradation and plant metal nutrition. Mol. Cell 69, 953– 964. https://doi.org/10.1016/j.molcel.2018.02.009

Faulkner, C., 2013. Receptor-mediated signaling at plasmodesmata. Front. Plant Sci. 4, 521. https://doi.org/10.3389/fpls.2013.00521

Faulkner, C., Petutschnig, E., Benitez-Alfonso, Y., Beck, M., Robatzek, S., Lipka, V., Maule, A.J., 2013. LYM2-dependent chitin perception limits molecular flux via plasmodesmata. Proc. Natl. Acad. Sci. U. S. A. 110, 9166–70. https://doi.org/10.1073/pnas.1203458110

Fernandez-Calvino, L., Faulkner, C., Walshaw, J., Saalbach, G., Bayer, E., Benitez-Alfonso, Y., Maule, A., 2011. Arabidopsis plasmodesmal proteome. PLoS One 6. https://doi.org/10.1371/journal.pone.0018880

Gallagher, K.L., Sozzani, R., Lee, C.-M., 2014. Intercellular Protein Movement: Deciphering the Language of Development. Annu. Rev. Cell Dev. Biol. 30, 207–233. https://doi.org/10.1146/annurev-cellbio-100913-012915

Gaus, K., 2014. ScienceDirect The organisation of the cell membrane?: do proteins rule lipids? ‘re’ mie Rossy, Yuanqing Ma and Katharina Gaus 54–59. https://doi.org/10.1016/j.cbpa.2014.04.009

Grison, M.S., Brocard, L., Fouillen, L., Nicolas, W., Wewer, V., Dörmann, P., Nacir, H., Benitez-Alfonso, Y., Claverol, S., Germain, V., Boutté, Y., Mongrand, S., Bayer, E.M., 2015. Specific membrane lipid composition is important for plasmodesmata function in Arabidopsis. Plant Cell 27, 1228–50. https://doi.org/10.1105/tpc.114.135731

Grison, M.S., Brocard, L., Fouillen, L., Nicolas, W., Wewer, V., Dörmann, P., Nacir, H., Benitez-Alfonso, Y., Claverol, S., Germain, V., Boutté, Y., Mongrand, S., Bayer, E.M., 2015. Specific membrane lipid composition is important for plasmodesmata function in arabidopsis. Plant Cell 27. https://doi.org/10.1105/tpc.114.135731

Gronnier, J., Crowet, J.-M., Habenstein, B., Nasir, M.N., Bayle, V., Hosy, E., Platre, M.P., Gouguet, P., Raffaele, S., Martinez, D., Grelard, A., Loquet, A., Simon-Plas, F., Gerbeau-Pissot, P., Der, C., Bayer, E.M., Jaillais, Y., Deleu, M., Germain, V., Lins, L., Mongrand, S., 2017. Structural basis for plant plasma membrane protein dynamics and organization into functional nanodomains. Elife 6. https://doi.org/10.7554/eLife.26404

Gronnier, J., Crowet, J.-M., Habenstein, B., Nasir, M.N., Bayle, V., Hosy, E., Platre, M.P., Gouguet, P., Raffaele, S., Martinez, D., Grelard, A., Loquet, A., Simon-Plas, F., Gerbeau-Pissot, P., Der, C., Bayer, E.M., Jaillais, Y., Deleu, M., Germain, V., Lins, L., Mongrand, S., 2017. Structural basis for plant plasma membrane protein dynamics and organization into functional nanodomains. Elife 6, 1–24. https://doi.org/10.7554/eLife.26404

Hartmann, M.A., Perret, A.M., Carde, J.P., Cassagne, C., Moreau, P., 2002. Inhibition of the sterol pathway in leek seedlings impairs phosphatidylserine and glucosylceramide synthesis but triggers an accumulation of triacylglycerols. Biochim. Biophys. Acta - Mol. Cell Biol. Lipids 1583, 285–296. https://doi.org/10.1016/S1388-1981(02)00249-4

He, J.-X., Fujioka, S., Li, T.-C., Kang, S.G., Seto, H., Takatsuto, S., Yoshida, S., Jang, J.-C., 2003. Sterols regulate development and gene expression in Arabidopsis. Plant Physiol. 131, 1258–1269. https://doi.org/10.1104/pp.014605.syndrome

Hem, S., Rofidal, V., Sommerer, N., Rossignol, M., 2007. Novel subsets of the Arabidopsis plasmalemma phosphoproteome identify phosphorylation sites in secondary active transporters. Biochem. Biophys. Res. Commun. 363, 375–380. https://doi.org/10.1016/j.bbrc.2007.08.177

Hofman, E.G., Ruonala, M.O., Bader, A.N., van den Heuvel, D., Voortman, J., Roovers, R.C., Verkleij, A.J., Gerritsen, H.C., van Bergen En Henegouwen, P.M.P., 2008. EGF induces coalescence of different lipid rafts. J. Cell Sci. 121, 2519–2528. https://doi.org/10.1242/jcs.028753

Hove, A. ten, Bochdanovits, Z., Jansweijer, V.M.A., Koning, F.G., Berke, L., Sanchez-Perez, G., Scheres, B., Heidstra, R., 2011. Probing the roles of LRR RLK genes in Arabidopsis thaliana roots using a custom T-DNA insertion set. Plant Mol Biol 76, 69–83. https://doi.org/10.1007/s11103-011-9769-x

Hsu, J.L., Wang, L.Y., Wang, S.Y., Lin, C.H., Ho, K.C., Shi, F.K., Chang, I.F., 2009. Functional phosphoproteomic profiling of phosphorylation sites in membrane fractions of salt-stressed Arabidopsis thaliana. Proteome Sci. 7, 42. https://doi.org/10.1186/1477-5956-7-42

Isner, J.C., Begum, A., Nuehse, T., Hetherington, A.M., Maathuis, F.J.M., 2018. KIN7 kinase regulates the vacuolar TPK1 K + channel during stomatal closure. Curr. Biol. 28, 466– 472. https://doi.org/10.1016/j.cub.2017.12.046

Jarsch, I.K., Konrad, S.S.A., Stratil, T.F., Urbanus, S.L., Szymanski, W., Braun, P., Braun, K.-H.H., Ott, T., 2014. Plasma Membranes Are Subcompartmentalized into a Plethora of Coexisting and Diverse Microdomains in Arabidopsis and Nicotiana benthamiana. Plant Cell 26, 1698–1711. https://doi.org/10.1105/tpc.114.124446

Keinath, N.F., Kierszniowska, S., Lorek, J., Bourdais, G., Kessler, S.A., Shimosato-Asano, H., Grossniklaus, U., Schulze, W.X., Robatzek, S., Panstruga, R., 2010. PAMP (Pathogen-associated Molecular Pattern)-induced changes in plasma membrane compartmentalization reveal novel components of plant immunity. J. Biol. Chem. 285, 39140–39149. https://doi.org/10.1074/jbc.M110.160531

Kierszniowska S, Seiwert B S.W., 2009. Definition of Arabidopsis sterol-rich membrane microdomains by differential treatment with methyl-beta-cyclodextrin and quantitative proteomics. Mol Cell Proteomics Apr;8(4):6.

Klepikova, A. V, Kasianov, A.S., Gerasimov, E.S., Logacheva, M.D., Penin, A.A., 2016. Ahigh resolution map of the Arabidopsis thaliana developmental transcriptome based on RNA-seq profiling 1058–1070. https://doi.org/10.1111/tpj.13312

Kline, K.G., Barrett-Wilt, G. a, Sussman, M.R., 2010. In planta changes in protein phosphorylation induced by the plant hormone abscisic acid. Proc. Natl. Acad. Sci. U. S. A. 107, 15986–15991. https://doi.org/10.1073/pnas.1007879107

Konrad, S.S.A., Popp, C., Stratil, T.F., Jarsch, I.K., Thallmair, V., Folgmann, J., Marín, M., Ott, T., 2014. S-acylation anchors remorin proteins to the plasma membrane but does not primarily determine their localization in membrane microdomains. New Phytol. 203, 758–769. https://doi.org/10.1111/nph.12867

Kragler, F., Monzer, J., Shash, K., Xoconostle-Cázares, B., Lucas, W.J., 1998. Cell-to-cell transport of proteins: Requirement for unfolding and characterization of binding to a putative plasmodesmal receptor. Plant J. 15, 367–381. https://doi.org/10.1046/j.1365-313X.1998.00219.x

Kumar, M., Yusuf, M.A., Yadav, P., Narayan, S., Kumar, M., Cushman, J.C., 2019. Overexpression of Chickpea defensin gene confers tolerance to water-deficit stress in Arabidopsis thaliana. Front. Plant Sci. 10, 290. https://doi.org/10.3389/fpls.2019.00290

Lee, E., Vanneste, S., Pérez-sancho, J., Benitez-Fuente, F., Strelau, M., Macho, A.P., Botella, M.A., Friml, J., Rosado, A., 2019. Ionic stress enhances ER – PM connectivity via site expansion in Arabidopsis. PNAS 116, 1420–1429. https://doi.org/10.1073/pnas.1818099116

Lee, J.-Y., Wang, X., Cui, W., Sager, R., Modla, S., Czymmek, K., Zybaliov, B., van Wijk, K., Zhang, C., Lu, H., Lakshmanan, V., 2011. A Plasmodesmata-Localized Protein Mediates Crosstalk between Cell-to-Cell Communication and Innate Immunity in Arabidopsis. Plant Cell Online 23, 3353–3373. https://doi.org/10.1105/tpc.111.087742

Levy, A., Erlanger, M., Rosenthal, M., Epel, B.L., 2007. A plasmodesmata-associated beta-1,3-glucanase in Arabidopsis. Plant J. 49, 669–682. https://doi.org/10.1111/j.1365-313X.2006.02986.x

Levy, A., Zheng, J.Y., Lazarowitz, S.G., 2015. Synaptotagmin SYTA Forms ER-Plasma Membrane Junctions that Are Recruited to Plasmodesmata for Plant Virus Movement. Curr. Biol. 25, 2018–2025. https://doi.org/10.1016/j.cub.2015.06.015

Lexy, R.O., Kasai, K., Clark, N., Fujiwara, T., Sozzani, R., Gallagher, K.L., 2018. Exposure to heavy metal stress triggers changes in plasmodesmatal permeability via deposition and breakdown of callose 69, 3715–3728. https://doi.org/10.1093/jxb/ery171

Lim, G.H., Shine, M.B., De Lorenzo, L., Yu, K., Cui, W., Navarre, D., Hunt, A.G., Lee, J.Y., Kachroo, A., Kachroo, P., 2016. Plasmodesmata Localizing Proteins Regulate Transport and Signaling during Systemic Acquired Immunity in Plants. Cell Host Microbe 19, 541–549. https://doi.org/10.1016/j.chom.2016.03.006

Liu, L., Liu, C., Hou, X., Xi, W., Shen, L., Tao, Z., Wang, Y., Yu, H., 2012. FTIP1 is an essential regulator required for florigen transport. PLoS Biol. 10. https://doi.org/10.1371/journal.pbio.1001313

MacGregor, D.R., Deak, K.I., Ingram, P.A., Malamy, J.E., 2008. Root system architecture in Arabidopsis grown in culture is regulated by sucrose uptake in the aerial tissues. Plant Cell 20, 2643–2660. https://doi.org/10.1105/tpc.107.055475

Martiniere, A., Lavagi, I., Nageswaran, G., Rolfe, D.J., Maneta-Peyret, L., Luu, D.-T., Botchway, S.W., Webb, S.E.D., Mongrand, S., Maurel, C., Martin-Fernandez, M.L., Kleine-Vehn, J., Friml, J., Moreau, P., Runions, J., 2012. Cell wall constrains lateral diffusion of plant plasma-membrane proteins. Proc. Natl. Acad. Sci. 109, 12805–12810. https://doi.org/10.1073/pnas.1202040109

Maule, A.J., Gaudioso-pedraza, R., Benitez-alfonso, Y., 2013. Callose deposition and symplastic connectivity are regulated prior to lateral root emergence. Commun. Intergrative Biol. 6:6, e26531.

Minami, A., Fujiwara, M., Furuto, A., Fukao, Y., Yamashita, T., Kamo, M., Kawamura, Y., Uemura, M., 2009. Alterations in detergent-resistant plasma membrane microdomains in Arabidopsis thaliana during cold acclimation. Plant Cell Physiol. 50, 341–359. https://doi.org/10.1093/pcp/pcn202

Miyashima, S., Roszak, P., Sevilem, I., Toyokura, K., Blob, B., Heo, J., Mellor, N., Helprinta-rahko, H., Otero, S., Smet, W., Boekschoten, M., Hooiveld, G., Hashimoto, K., Smetana, O., Siligato, R., Wallner, E., Mähönen, A.P., Kondo, Y., 2019. Mobile PEAR transcription factors integrate positional cues to prime cambial growth. Nature 565, 490– 494. https://doi.org/10.1038/s41586-018-0839-y

Morsomme, P., Dambly, S., Maudoux, O., Boutry, M., 1998. Single point mutations distributed in 10 soluble and membrane regions of the Nicotiana plumbaginifolia plasma membrane PMA2 H+-ATPase activate the enzyme and modify the structure of the Cterminal region. J. Biol. Chem. 273, 34837–34842. https://doi.org/10.1074/jbc.273.52.34837

Nicolas, W.J., Grison, M.S., Bayer, E.M., 2017. Shaping intercellular channels of plasmodesmata: the structure-to-function missing link. J. Exp. Bot. https://doi.org/10.1093/jxb/erx225

Niittylä, T., Fuglsang, A.T., Palmgren, M.G., Frommer, W.B., Schulze, W.X., 2007. Temporal analysis of sucrose-induced phosphorylation changes in plasma membrane proteins of Arabidopsis. Mol. Cell. Proteomics 6, 1711–1726. https://doi.org/10.1074/mcp.M700164-MCP200

Nikonorova, N., Broeck, L. Van Den, Zhu, S., Cotte, B. Van De, 2018. Early mannitol-triggered changes in the Arabidopsis leaf (phospho) proteome reveal growth regulators 69, 4591–4607. https://doi.org/10.1093/jxb/ery261

Otero, S., Helariutta, Y., Benitez-Alfonso, Y., 2016. Symplastic communication in organ formation and tissue patterning. Curr. Opin. Plant Biol. 29, 21–28. https://doi.org/10.1016/j.pbi.2015.10.007

Péret, B., Li, G., Zhao, J., Band, L.R., Voß, U., Postaire, O., Luu, D.-T., Da Ines, O., Casimiro, I., Lucas, M., Wells, D.M., Lazzerini, L., Nacry, P., King, J.R., Jensen, O.E., Schäffner, A.R., Maurel, C., Bennett, M.J., 2012. Auxin regulates aquaporin function to facilitate lateral root emergence. Nat. Cell Biol. 14, 991–8. https://doi.org/10.1038/ncb2573

Perraki, A., Gronnier, J., Gouguet, P., Boudsocq, M., Deroubaix, A.-F., Simon, V., German-Retana, S., Legrand, A., Habenstein, B., Zipfel, C., Bayer, E., Mongrand, S., Germain, V., 2018. REM1.3’s phospho-status defines its plasma membrane nanodomain organization and activity in restricting PVX cell-to-cell movement. PLoS Pathog. 14(11): e1.

Prak, S., Hem, S., Boudet, J., Viennois, G., Sommerer, N., Rossignol, M., Maurel, C., Santoni, V., 2008. Multiple Phosphorylations in the C-terminal Tail of Plant Plasma Membrane Aquaporins. Mol. Cell. Proteomics 7, 1019–1030. https://doi.org/10.1074/mcp.M700566-MCP200

Raffaele, S., Mongrand, S., Gamas, P., Niebel, A., Ott, T., 2007. Genome-Wide Annotation of Remorins, a Plant-Specific Protein Family: Evolutionary and Functional Perspectives. Plant Physiol. 145, 593–600. https://doi.org/10.1104/pp.107.108639

Reagan, B.C., Ganusova, E.E., Fernandez, J.C., Mccray, T.N., 2018. RNA on the move?: the plasmodesmata perspective. Plant Sci. 275, 1–10. https://doi.org/10.1016/j.plantsci.2018.07.001

Roycewicz, P., Malamy, J.E., 2012. Dissecting the effects of nitrate, sucrose and osmotic potential on Arabidopsis root and shoot system growth in laboratory assays. Phil. Trans. R. Soc. B 367, 1489–1500. https://doi.org/10.1098/rstb.2011.0230

Salmon, M.S., Bayer, E.M., 2013. Dissecting plasmodesmata molecular composition by mass spectrometry-based proteomics. Front. Plant Sci. 3, 307. https://doi.org/10.3389/fpls.2012.00307

Shahollari, B., Peskan-Berghöfer, T., Oelmüller, R., 2004. Receptor kinases with leucine-rich repeats are enriched in Triton X-100 insoluble plasma membrane microdomains from plants. Physiol. Plant. 122, 397–403. https://doi.org/10.1111/j.1399-3054.2004.00414.x

Shahollari, B., Varma, A., Oelmüller, R., 2005. Expression of a receptor kinase in Arabidopsis roots is stimulated by the basidiomycete Piriformospora indica and the protein accumulates in Triton X-100 insoluble plasma membrane microdomains. J. Plant Physiol. 162, 945–958. https://doi.org/10.1016/j.jplph.2004.08.012

Simpson, C., Thomas, C., Findlay, K., Bayer, E., Maule, A.J., 2009. An Arabidopsis GPI-anchor plasmodesmal neck protein with callose binding activity and potential to regulate cell-to-cell trafficking. Plant Cell 21, 581–594. https://doi.org/10.1105/tpc.108.060145

Sivaguru, M., Fujiwara, T., Samaj, J., Baluska, F., Yang, Z., Osawa, H., Maeda, T., Mori, T., Volkmann, D., Matsumoto, H., 2000. Aluminum-induced 1-->3-beta-D-glucan inhibits cell-to-cell trafficking of molecules through plasmodesmata. A new mechanism of aluminum toxicity in plants. Plant Physiol. 124, 991–1006. https://doi.org/10.1104/pp.124.3.991

Srivastava, V., Malm, E., Sundqvist, G., Bulone, V., 2013. Quantitative proteomics reveals that plasma membrane microdomains from poplar cell suspension cultures are enriched in markers of signal transduction, molecular transport, and callose biosynthesis *. Mol. Cell. Proteomics 12. 12, 3874–3885. https://doi.org/10.1074/mcp.M113.029033

Stahl, Y., Faulkner, C., 2015. Receptor Complex Mediated Regulation of Symplastic Traffic. Trends Plant Sci. xx, 1–10. https://doi.org/10.1016/j.tplants.2015.11.002

Stahl, Y., Grabowski, S., Bleckmann, A., Kühnemuth, R., Weidtkamp-Peters, S., Pinto, K.G., Kirschner, G.K., Schmid, J.B., Wink, R.H., Hülsewede, A., Felekyan, S., Seidel, C.A.M., Simon, R., 2013. Moderation of arabidopsis root stemness by CLAVATA1 and ARABIDOPSIS CRINKLY4 receptor kinase complexes. Curr. Biol. 23, 362–371. https://doi.org/10.1016/j.cub.2013.01.045

Stahl, Y., Simon, R., 2013. Gated communities: Apoplastic and symplastic signals converge at plasmodesmata to control cell fates. J. Exp. Bot. 64, 5237–5241. https://doi.org/10.1093/jxb/ert245

Szymanski, W.G., Zauber, H., Erban, A., Wu, X.N., Schulze, W.X., 2015. Cytoskeletal components define protein location to membrane microdomains. Mol. Cell. Proteomics M114.046904-. https://doi.org/10.1074/mcp.M114.046904

Thomas, C.L., Bayer, E.M., Ritzenthaler, C., Fernandez-Calvino, L., Maule, A.J., 2008. Specific targeting of a plasmodesmal protein affecting cell-to-cell communication. PLoS Biol. 6. https://doi.org/10.1371/journal.pbio.0060007

Thomas, C.L., Bayer, E.M., Ritzenthaler, C., Fernandez-Calvino, L., Maule, A.J., 2008. Specific targeting of a plasmodesmal protein affecting cell-to-cell communication. PLoS Biol. 6, 0180–0190. https://doi.org/10.1371/journal.pbio.0060007

Tilsner, J., Amari, K., Torrance, L., 2011. Plasmodesmata viewed as specialised membrane adhesion sites. Protoplasma 248, 39–60. https://doi.org/10.1007/s00709-010-0217-6

Tilsner, J., Nicolas, W., Rosado, A., Bayer, E.M., 2016. Staying tight: plasmodesmata membrane contact sites and the control of cell-to-cell connectivity. Annu. Rev. Plant Biol. 67, 337–64.

Tylewicz, S., Bhalerao, R.P., 2018. Photoperiodic control of seasonal growth is mediated by ABA acting on cell-cell communication. Science (80-.). 8576, 1–9. https://doi.org/10.1126/science.aan8576

Vaddepalli, P., Herrmann, A., Fulton, L., Oelschner, M., Hillmer, S., Stratil, T.F., Fastner, A., Hammes, U.Z., Ott, T., Robinson, D.G., Schneitz, K., 2014. The C2-domain protein QUIRKY and the receptor-like kinase STRUBBELIG localize to plasmodesmata and mediate tissue morphogenesis in Arabidopsis thaliana. Development 141, 4139–4148. https://doi.org/10.1242/dev.113878

Vaten, A., Dettmer, J., Wu, S., Stierhof, Y.D., Miyashima, S., Yadav, S.R., Roberts, C.J., Campilho, A., Bulone, V., Lichtenberger, R., Lehesranta, S., Mähönen, A.P., Kim, J.Y., Jokitalo, E., Sauer, N., Scheres, B., Nakajima, K., Carlsbecker, A., Gallagher, K.L., Helariutta, Y., 2011. Callose Biosynthesis Regulates Symplastic Trafficking during Root Development. Dev. Cell 21, 1144–1155. https://doi.org/10.1016/j.devcel.2011.10.006

Wang, X., Sager, R., Cui, W., Zhang, C., Lu, H., Lee, J., 2013. Salicylic acid regulates Plasmodesmata closure during innate immune responses in Arabidopsis. Plant Cell 25, 2315–29. https://doi.org/10.1105/tpc.113.110676

Wattelet-Boyer, V., Brocard, L., Jonsson, K., Esnay, N., Joubès, J., Domergue, F., Mongrand, S., Raikhel, N., Bhalerao, R.P., Moreau, P., Boutté, Y., 2016. Enrichment of hydroxylated C24-and C26-acyl-chain sphingolipids mediates PIN2 apical sorting at trans-Golgi network subdomains. Nat. Commun. 7, 12788. https://doi.org/10.1038/ncomms12788

Wu, S., O’Lexy, R., Xu, M., Sang, Y., Chen, X., Yu, Q., Gallagher, K.L., 2016. Symplastic signaling instructs cell division, cell expansion, and cell polarity in the ground tissue of Arabidopsis thaliana roots. Proc. Natl. Acad. Sci. U. S. A. 113, 11621–11626. https://doi.org/10.1073/pnas.1610358113

Wu, X.N., Sanchez Rodriguez, C., Pertl-Obermeyer, H., Obermeyer, G., Schulze, W.X., Wu XN, Sanchez Rodriguez C, Pertl-Obermeyer H, Obermeyer G S.W., 2013. Sucrose-induced receptor kinase SIRK1 regulates a plasma membrane aquaporin in Arabidopsis. Mol Cell Proteomics. 12, 2856–73. https://doi.org/10.1074/mcp.M113.029579

Xu, B., Cheval, C., Laohavisit, A., Hocking, B., Chiasson, D., Olsson, T.S.G., Shirasu, K., Faulkner, C., Gilliham, M., 2017. A calmodulin-like protein regulates plasmodesmal closure during bacterial immune responses. New Phytol. 215, 77–84. https://doi.org/10.1111/nph.14599

Xue, L., Wang, P., Wang, L., Renzi, E., Radivojac, P., Tang, H., Arnold, R., Zhu, J., Tao, W.A., 2013. Quantitative Measurement of Phosphoproteome Response to Osmotic Stress in Arabidopsis Based on Library-Assisted eXtracted Ion Chromatogram (LAXIC)* ?. Mol. Cell. Proteomics 12.8, 2354–2369. https://doi.org/10.1074/mcp.O113.027284

Zavaliev, R., Ueki, S., Epel, B.L., Citovsky, V., 2011. Biology of callose (β-1,3-glucan) turnover at plasmodesmata. Protoplasma 248, 117–130. https://doi.org/10.1007/s00709-010-0247-0

Zhou, A., Ma, H., Fen, S., Gong, S., Wang, J., 2018. A Novel Sugar Transporter from Affects Sugar Metabolism and Confers Osmotic and Oxidative Stress Tolerance in Arabidopsis. Int. J. Mol. Sci. 19, 1–10. https://doi.org/10.3390/ijms19020497

